# Stress-induced organismal death is genetically regulated by the mTOR-Zeste-Phae1 axis

**DOI:** 10.1101/2024.12.25.630336

**Authors:** Takashi Matsumura, Masasuke Ryuda, Hitoshi Matsumoto, Takumi Kamiyama, Shu Kondo, Akira Nakamura, Yoichi Hayakawa, Ryusuke Niwa

## Abstract

All organisms are exposed to various stressors, which can sometimes lead to organismal death, depending on their intensity. While stress-induced organismal death has been observed in many species, the underlying mechanisms remain unclear. In this study, we investigated the molecular mechanisms of stress-induced organismal death in the fruit fly *Drosophila melanogaster*. We identified a chymotrypsin-like serine protease *Phaedra1* (*Phae1*) as a death mediator in *D. melanogaster* larvae. *Phae1* expression was upregulated by lethal heat stress (40 °C) but not non-lethal heat stress (38 °C or lower). The most prominent induction of *Phae1* occurred in the central nervous system (CNS). We found neuro-specific knockdown of *Phae1* increased survival and reduced neuronal caspase activity following exposure to lethal heat stress, suggesting that the transcriptional upregulation of *Phae1* in the CNS is essential for stress-induced organismal death. We next found via bioinformatic and biochemical analyses that the transcription factor Zeste (Z) bound the *Phae1* enhancer region and that *z* loss-of-function impaired *Phae1* induction in the CNS, increasing survival following lethal heat stress. In addition, we found via chemical screening that rapamycin, a chemical inhibitor of mechanistic target of rapamycin (mTOR), suppressed *Phae1* expression. Neuro-specific knockdown of *mTor* reduced the protein levels of both Phae1 and Z, leading to an increase in survival following lethal heat stress. Together, these results indicate that heat stress-induced organismal death in *D. melanogaster* larvae is regulated by a genetically encoded transcriptional signaling pathway involving the mTOR-Z-Phae1 axis.

## Introduction

In nature, living organisms must survive various stressors, such as low and high temperatures, dehydration, ultraviolet radiation, infection, fighting, crowding, and lack of food (1–8). Stress affects organismal physiology in part by inducing the expression of stress-responsive genes related to specific metabolic pathways, autophagy, and cell death (9–12). Organisms can survive stress levels that remain below their species-specific thresholds (13, 14), but when stress exceeds these thresholds, it can lead to organismal death (15). Stress-induced organismal death occurs in various animals, including humans (16, 17), often resulting from lethal stress exposure without any visible external damage (6). It remains unclear whether stress-induced organismal death is merely a secondary consequence of some sort of physiological breakdown or an active process driven by genetic mechanisms triggered by the excessive stress. A few pioneering studies using the nematode *Caenorhabditis elegans* suggested the involvement of genetic mechanisms in stress-induced organismal death. For example, nematodes exposed to lethal heat stress exhibit activation of the protease cascade along with a specific transcriptional pathway that promotes abnormal cell death (18, 19). Despite these studies, however, the key mediators underlying stress-induced organismal death remain largely unknown.

The fruit fly *Drosophila melanogaster* has long been a valuable model for studying the molecular mechanisms of animal physiology (20). In recent years, *D. melanogaster* has also emerged as an important model for investigating stress-induced injury pathways (21). To uncover the mechanisms underlying stress-induced organismal death, we first wanted to identify a reliable hallmark of organismal death. Such a hallmark would hopefully allow us to trace the origin of organismal death to a specific organ or tissue and then investigate the molecular regulatory events occurring there. In this study, we used an RNA-seq approach to identify the *Drosophila* gene *Phaedral1* (*Phae1*), which exhibits stress-specific transcriptional activation. We also examined the tissues where *Phae1* is expressed and the mechanism underlying *Phae1* expression. In stressed larvae, *Phae1* is prominently expressed in the central nervous system (CNS). Lethal heat stress activates *Phae1* expression via the transcription factor Zeste (Z). Furthermore, we found this Z-dependent induction of *Phae1* expression in response to lethal heat stress is regulated by the mechanistic target of rapamycin (mTOR). We were able to confirm that this mTOR-Z-Phae1 pathway is essential for heat stress-induced organismal death, indicating that, at least in *D. melanogaster*, some forms of stress-induced organismal death are regulated by genetically-encoded transcriptional or signal transduction relays.

## Results

### *Phaedra1 (Phae1)*, encoding a serine protease, is a lethal stress-responsive gene

Before investigating the molecular mechanism underlying stress-induced organismal death, we examined the survival of *D. melanogaster* larvae exposed to 30 minutes of heat stress at various temperatures. No larvae died from exposure to temperatures of 38 °C or lower, and only a few succumbed to a 30-min exposure to 39 °C. In contrast, almost all larvae died from a 30-min exposure to 40 °C, indicating that there is only a narrow 2 °C window between survivable and lethal stress (Fig.1A). We therefore used these two temperatures as non-lethal (38 °C) and lethal (40 °C) stressors. To identify genes involved in mediating heat stress-induced death, we looked for genes induced specifically by lethal stress in the larval fat body by performing a messenger RNA sequencing (mRNA-seq) transcriptome analysis comparing the response to non-lethal and lethal heat stress. We identified 78 genes upregulated in larvae exposed to lethal heat stress versus those exposed to non-lethal heat stress (fold change > 2, FDR < 0.000001). These 78 candidate genes included proteolysis-related genes (10 genes) and chitin metabolism-related genes (14 genes) (SI Appendix, Fig. S1A, Dataset S01). We next confirmed the specific lethal stress-induced upregulation of 23 of these candidate genes by quantitative reverse transcription PCR (qPCR) (SI Appendix, Fig. S1A, Dataset S01).

**Figure 1.**
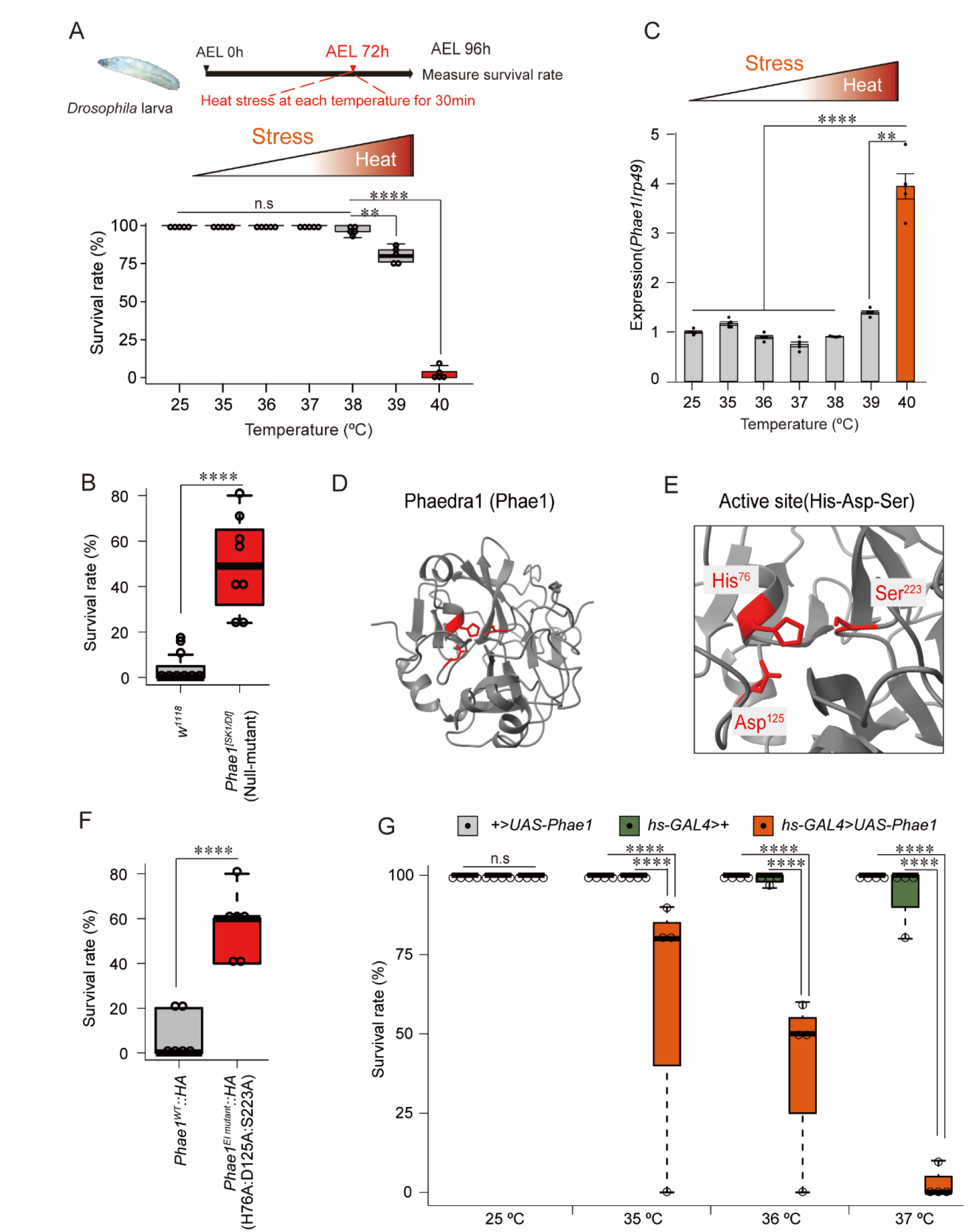
*Phae1* is a lethal heat stress-responsive death mediator gene. (A) *Drosophila* larval survival following 30 min of heat stress at the indicated temperatures. \*\**P* < 0.01, \*\*\*\**P* < 0.001 (Fisher’s exact test). N = 5 independent technical replicates, n = 25 independent biological replicates. (B) Control (*w^1118^*) and *Phae1* mutant larval survival following a 30-min exposure to lethal heat stress at 40 °C. All values are means ± SE. \*\*\*\**P* < 0.001 (Fisher’s exact test). N = 8 independent technical replicates, n = 25 independent biological replicates. (C) *Phae1* mRNA levels after a 30-min heat stress exposure at the indicated temperatures. \*\**P* < 0.01, \*\*\*\**P* < 0.001 (one-way ANOVA followed by Tukey’s HSD). N = 4 independent technical replicates. (D) Phae1 protein 3D structure. (E) Phae1 protease catalytic resides (Red area, His-Asp-Ser). (F) Survival of larvae with or without Phae1 protease function after exposure to lethal heat stress. \*\*\*\**P* < 0.001 (Fisher’s exact test). N = 6 independent technical replicates, n = 25 independent biological replicates. (G) Survival of larvae with or without *Phae1* overexpression following a 30-min exposure to the indicated temperature. All values are means ± SE. \*\*\*\**P* < 0.001 (Fisher’s exact test). N = 4 independent technical replicates, n = 25 independent biological replicates.

To screen these candidate mediator genes of stress-induced organismal death, we next exposed larvae with *in vivo* RNAi (*cg-GAL4>UAS-RNAi*) to lethal heat stress at 40 °C and measured their survival. In this screen, we found that knockdown of *Phaedra1* (*Phae1*), which encodes a chymotrypsin-like serine protease (22), increased larval survival (SI Appendix, Fig. S1B). This makes *Phae1* a candidate death mediator gene. We confirmed that a *Phae1* genetic null mutant (*Phae1^[SK1]/[Df]^*) phenocopied the *Phae1* RNAi result, with more mutant larvae surviving exposure to lethal heat stress than *w^1118^* control larvae (Fig. 1B).

We also found that lethal (40 °C) but not nonlethal (38 °C) heat stress increased the expression of *Phae1* (Fig.1C). After establishing a *Phae1-GAL4>UAS-GFP* (*Phae1>GFP*) transgenic reporter strain to visualize *Phae1* expression *in vivo*, we found increased *Phae1* expression upon exposure to various stressors, including low and high temperatures, the insecticides imidacloprid, methamidophos, and rotenone, as well as desiccation, but not starvation (SI Appendix Fig. S2).

We next wondered whether Phae1 protease activity is responsible for stress-induced organismal death. Phae1 protein is predicted to have a typical serine protease active site, consisting of Histidine^76^, Aspartic acid^125^, and Serine^223^ (His^76^, Asp^125^, Ser^223^) (23) (Fig. 1D and E). Thus, we established a knock-in *Phae1* strain with site-directed substitutions of these three amino acids to alanine (H76A:D125A:S223A) using the CRISPR-Cas9 technique (24). We chose alanine substitutions to minimize unfavorable steric effects (25). We used a single guide RNA (sgRNA) to replace the *Phae1* CDS, resulting in HA-tagged Phae1 proteins with or without the enzymatically inactive (EI) mutations in the Phae1 active site (*Phae1^WT^::HA* or *Phae1^EI^ ^mutant^::HA*). As expected, more EI mutant (*Phae1^EI^ ^mutant^::HA*) larvae than wild-type larvae (*Phae1^WT^::HA*) survived following exposure to lethal stress, suggesting Phae1 protease activity is critical for stress-induced organismal death (Fig. 1F).

We next asked whether forced overexpression of a *Phae1* transgene could enhance stress-induced organismal death. When we overexpressed *Phae1* in 3rd instar larvae (*hs-GAL4>UAS-Phae1*), we observed a reduction in their survival compared with control larvae lacking *Phae1* over-expression (*+>UAS-Phae1* and *hs-GAL4>+*) at each temperature; 25% of the larvae died at 35 °C, 50% of the larvae died at 36 °C, and all of the larvae died at 37 °C (Fig. 1G). These results are consistent with the hypothesis that *Phae1* promotes heat stress-induced organismal death.

### Neuronal *Phae1* affects survival following lethal stress

Using public single-cell RNA-seq data in *Flybase* (http://flybase.org/reports/FBgn0263234.htm) (26), we found *Phae1* is strongly expressed in several other larval tissues in addition to the fat body, including epidermal cells and the gut. We therefore used qPCR to determine whether *Phae1* expression is also induced in these other tissues upon exposure to lethal heat stress. Compared to control conditions at 25 °C and non-lethal stress at 38 °C, the lethal stress condition at 40 °C led to an upregulation of *Phae1* expression in various tissues (Fig. 2A). After calculating the fold change in *Phae1* expression for each tissue following lethal heat stress, we found the largest increase in the central nervous system (CNS) (Fig. 2B).

**Figure 2.**
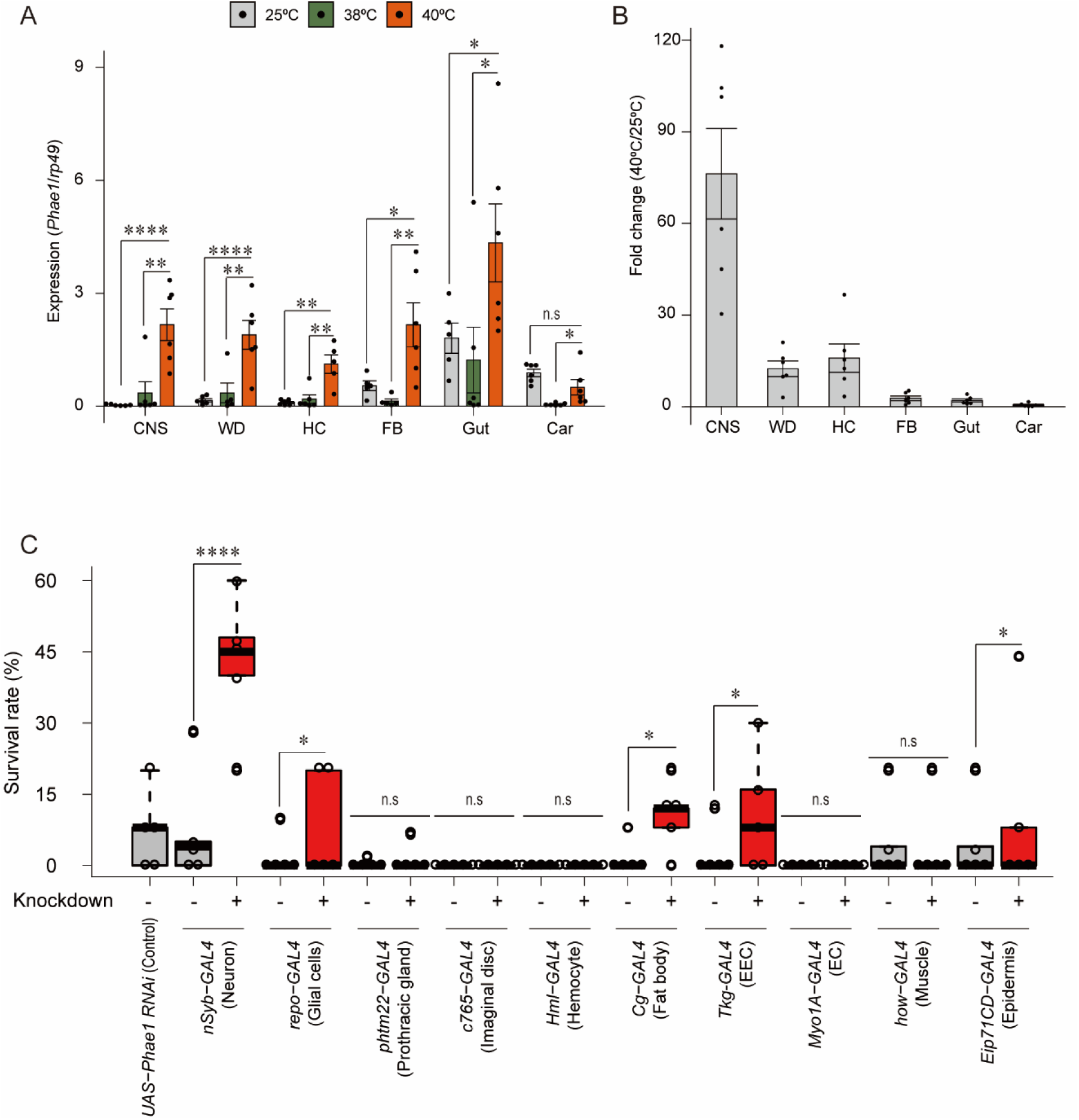
Neuronal *Phae1* expression is specifically upregulated by heat stress. (A) *Phae1* mRNA level in various tissues CNS; central nervous system, WD; wing disc, Hc; hemocyte, FB; fat boy, Gut, Car; carcass. \**P* < 0.05, \*\**P* < 0.01, \*\*\*\**P* < 0.001 (one-way ANOVA followed by Tukey’s HSD). N = 6 independent technical replicates. (B) *Phae1* mRNA fold change in various tissues. (C) Survival of larvae with or without tissue-specific *Phae1* knockdown following a 30-min exposure to 40 °C (EEC; enteroendocrine cells, EC; enterocyte,). All values are means ± SE. \**P* < 0.05, \*\**P* < 0.01, \*\*\*\**P* < 0.001 (Fisher’s exact test). N = 6 independent technical replicates, n = 25 independent biological replicates.

To clarify the importance of this increase in neuronal *Phae1* expression, we used the GAL4-UAS system (20) to investigate the effects of tissue-specific *Phae1* knockdown on larval survival following lethal heat stress. We found a dramatic increase in survival upon neuron-specific *Phae1* knockdown, slight increases with tissue-specific knockdown in the epidermis, enteroendocrine cells (EEC), enterocyte (EC), fat body, and glial cells, and no change with other tissue-specific drivers compared to control larvae (Fig. 2C, SI Appendix, Fig. S3A). We were also able to confirm with a second neuronal driver, *elav-GAL4*, that neuro-specific *Phae1* knockdown increased larval survival following lethal heat stress (SI Appendix, Fig. S3B). These results suggest neuronal *Phae1* is the most important for stress-induced organismal death.

Although *Phae1* is similar in sequence with it paralogue *Phaedra2* (*Phae2*), which also encodes a chymotrypsin-type serine protease, neuro-specific *Phae2* knockdown did not increase larval survival toward lethal heat stress (SI Appendix Fig. S3A). Thus, we focused only on *Phae1* for the rest of this study.

### *Phae1* promotes neuronal cell death along with caspase activation

We next wondered how neuronal Phae1 promotes stress-induced organismal death. Stress exposure sometimes induces apoptosis-like cell death in dying animals (27, 28), which inspired us to ask whether stress-induced cell death could be part of the mechanism underlying stress-induced organismal death. We first asked whether stress-induced cell death occurs in the *D. melanogaster* larval CNS after exposure to lethal heat stress and whether *Phae1* is involved in the induction of stress-dependent cell death. In the CNS of control larvae (*w^1118^*), we observed a massive induction of cell death by lethal stress at 40 °C, but not by non-lethal stress at 38 °C (Fig. 3A-C). *Phae1* knockout, however, reduced the number of TUNEL-positive dead cells (TUNEL^+^ cells) following lethal stress (Fig. 3C’, 3D-F, 3M), suggesting *Phae1* regulates stress-induced cell death in the larval CNS.

**Figure 3.**
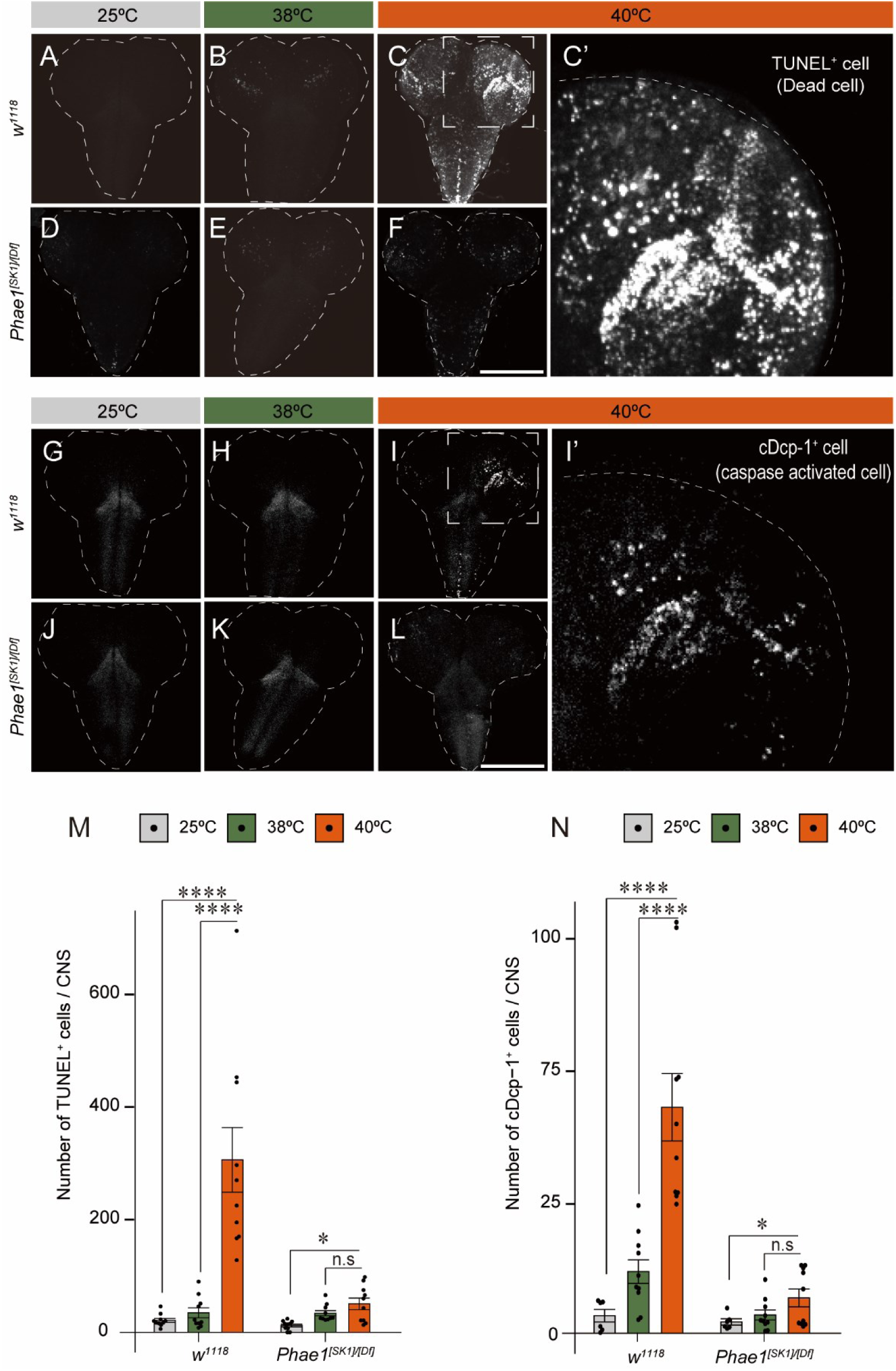
*Phae1* is associated with cell death induction and caspase activation. (A-F) TUNEL-stained CNS from *Drosophila* larvae exposed or unexposed to heat stress. Dotted lines encircle the larval CNS. Scale bar: 200 μm. (C’) Magnified view of the CNS of control (*w^1118^*) larvae following exposure to lethal heat stress. (G-I) CNS of larvae either exposed or unexposed to heat stress stained with a cleaved Dcp-1 (cDcp-1) antibody. Dotted lines encircle the larval CNS. Scale bar: 200 μm. (I’) Magnified view of the CNS of control (*w^1118^*) larvae following exposure to lethal heat stress. (M) Quantification of TUNEL-positive cells. (N) Quantification of cDcp-1-positive cells. All values are means ± SE. \**P* < 0.05, \*\*\*\**P* < 0.001 (one-way ANOVA followed by Tukey’s HSD). N = 8 independent technical replicates.

To determine whether Phae1 promotes cell death by regulating active caspase levels, we next quantified the levels of the active form of the *D. melanogaster* caspase 3 homolog Dcp-1 (cDcp-1) between control and *Phae1* knockout larvae. Caspases are cysteine proteases associated with programmed cell death (29). Consistent with our earlier results, although we observed an increase in cDcp-1-positive cells (cDcp-1^+^ cells) in control larval brains following lethal heat stress (Fig.3G-I, 3I’), *Phae1* knockout prevented this stress-induced Dcp-1 activation (Fig. 3J-L, 3N). Neuron-specific *Phae1* knockdown inhibited both Dcp1-activation and cell death induction (SI Appendix, Fig. S4). We also found neuron-specific overexpression of the caspase inhibitor *p35* increased larval survival following lethal heat stress (SI Appendix, Fig. S5).

These results suggest Phae1-induced cell death, along with neuronal caspase activation, are responsible for heat stress-induced organismal death.

### The transcription factor Zeste (Z) regulates neuronal *Phae1* expression following lethal heat stress

Thus far, we have demonstrated a specific upregulation of *Phae1* expression following lethal but not non-lethal heat stress, implying the existence of lethal stress-specific transcriptional regulation. To identify such a transcription factor regulating *Phae1* expression, we used a luciferase-based reporter assay to analyze the *cis*-elements of the *Phae1* promoter. First, we created luciferase reporter vectors containing various genomic regions upstream of the *Phae1* gene. After transfecting these vectors into cultured *D. melanogaster* Schneider 2 (S2) cells, we measured *Phae1* promoter-driven luciferase activity following lethal heat stress at 42 °C for 30-min according to the previously described procedure as follows (30). Heat stress triggered *luciferase* expression from the reporter vector containing the 550 base pairs (bp) upstream of *Phae1* (*Phae1>Luc*). With enhancers of 540 bp or less, however, we did not observe any response to stress (Fig. 4A), suggesting the presence of a stress response element within the 10-bp span from 540 to 550 bp upstream of the *Phae1* coding sequence. When we used this region to search the JASPAR transcription factor binding profile database (31), we found a predicted binding site for Zeste (Z) (Fig. 4A, SI Appendix, Fig. S6). Z is a trihelix transcription factor (32) with a helix-turn-helix type DNA binding domain (SI Appendix, Fig. S7A) (33).

**Figure 4.**
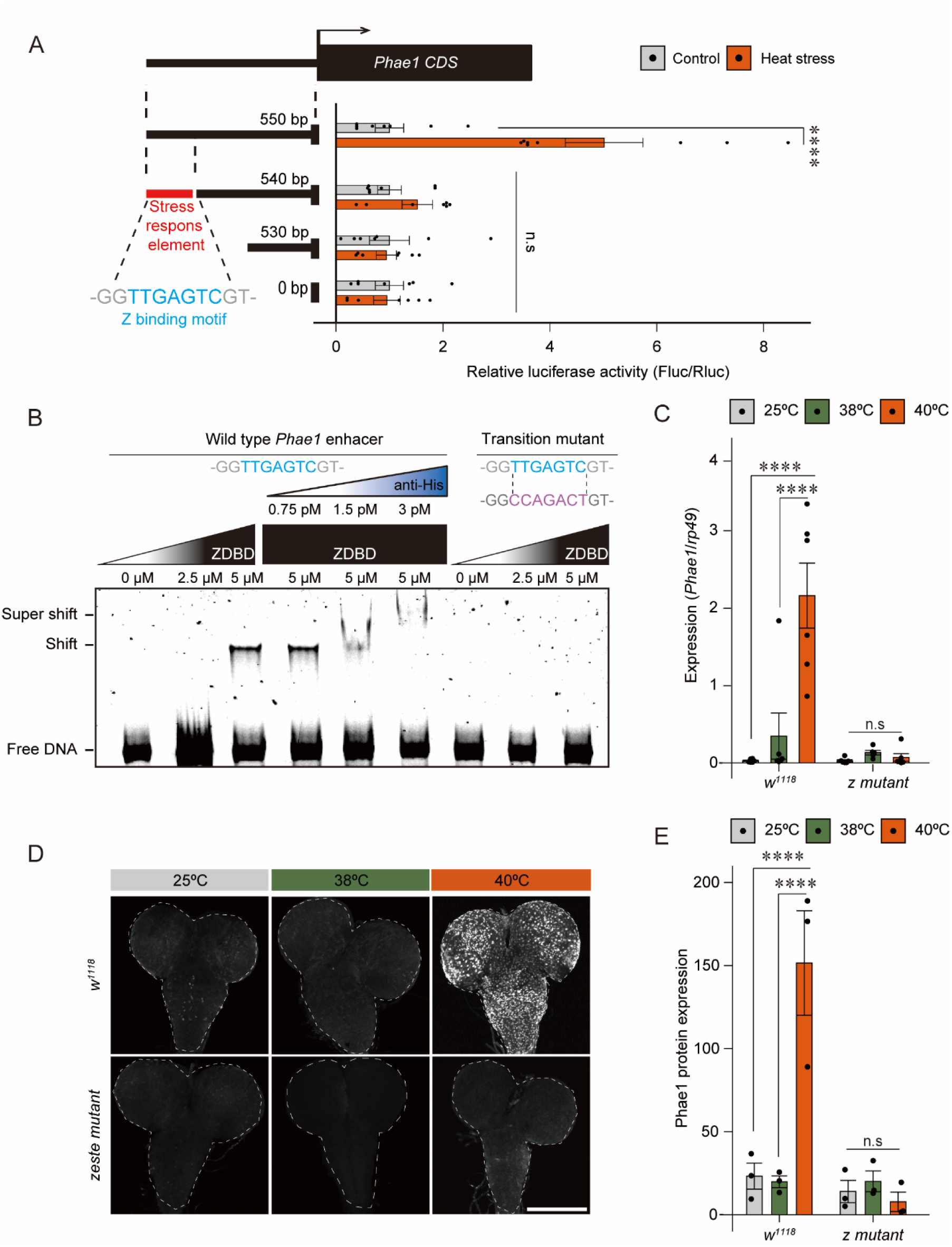
The Z transcription factor regulates *Phae1* expression following heat stress exposure. (A) Dual luciferase assay for identifying transcription factors that regulate *Phae1* gene expression following heat stress exposure at 42 °C for 30-min. \*\*\*\**P* < 0.001 (one-way ANOVA followed by Tukey’s HSD). N = 8 independent technical replicates. (B) Gel shift assay confirming that Z protein binds the *Phae1* enhancer. N = 3 independent technical replicates. (C) *Phae1* mRNA levels in control (*w^1118^*) and *z* mutant larvae. \*\*\*\**P* < 0.001 (one-way ANOVA followed by Tukey’s HSD). (D) Anti-Phae1 staining of the CNS of control (*w^111^*) and z mutant larvae exposed or unexposed to heat stress. Dotted lines encircle the larval CNS. Scale bar: 200 μm. (E) Quantification of Phae1 protein level. \*\*\*\**P* < 0.001 (one-way ANOVA followed by Tukey’s HSD). N = 3 independent technical replicates.

To determine whether Z protein binds the *Phae1* enhancer region, we performed an electrophoretic mobility shift assay (EMSA). The DNA fragment containing the wild-type *Phae1* enhancer region formed a complex with recombinant Z protein (ZDBD, His-tagged protein), resulting in a band shift (Fig. 4B). The addition of anti-His tag antibody induced a super shift due to increased molecular weight, suggesting the interaction of the antibody with a DNA-Z complex. This apparent DNA-Z protein interaction was completely abolished by mutation of the Z binding motif within the *Phae1* enhancer region, confirming the direct binding of Z to the *Phae1* enhancer region. We also found that *z* mutation reduced *Phae1* gene expression following lethal heat stress (Fig. 4C, SI Appendix, Fig. S8). We next examined neuronal Phae1 protein levels in control (*w^1118^*) and *z* mutant (*z^v77h^*) larvae. Although we observed the typical increase in Phae1 protein in the CNS of control larvae exposed to lethal heat stress, this increase was abolished in the CNS of *z* mutant larvae (Fig. 4D and 4E).

To further analyze Z protein *in vivo*, we used the CRISPR-Cas9 technique to produce a knock-in strain that added an N-terminal Ty1 tag (Z-Ty1) to the endogenous Z protein. Consistent with what we observed with *z* mRNA expression, we used this strain to confirm that lethal heat stress also increased neuronal Z protein levels (Fig. 5B, SI Appendix, Fig. S9). We then performed an *in vivo* chromatin immunoprecipitation (ChIP) analysis with an anti-Ty1 antibody and nuclear extracts from the Z-Ty1 larval CNS. In this analysis, we found evidence that anti-Ty1 increased the enrichment of the *Phae1* promoter region following lethal heat stress compared to control mouse IgG (Fig. 5C). We also found that both whole body knockout of *z* (Fig. 5D) and neuron-specific knockdown of *z* (Fig. 5E) increased the survival of larvae exposed to lethal heat stress. Both neuronal overexpression of either *z* or *Phae1*, however, completely abolished this z mutation-induced increase in larval survival of heat stress (SI Appendix, Fig. S10). We therefore concluded that Z is a crucial transcription factor for inducing *Phae1* expression after lethal heat stress.

**Figure 5.**
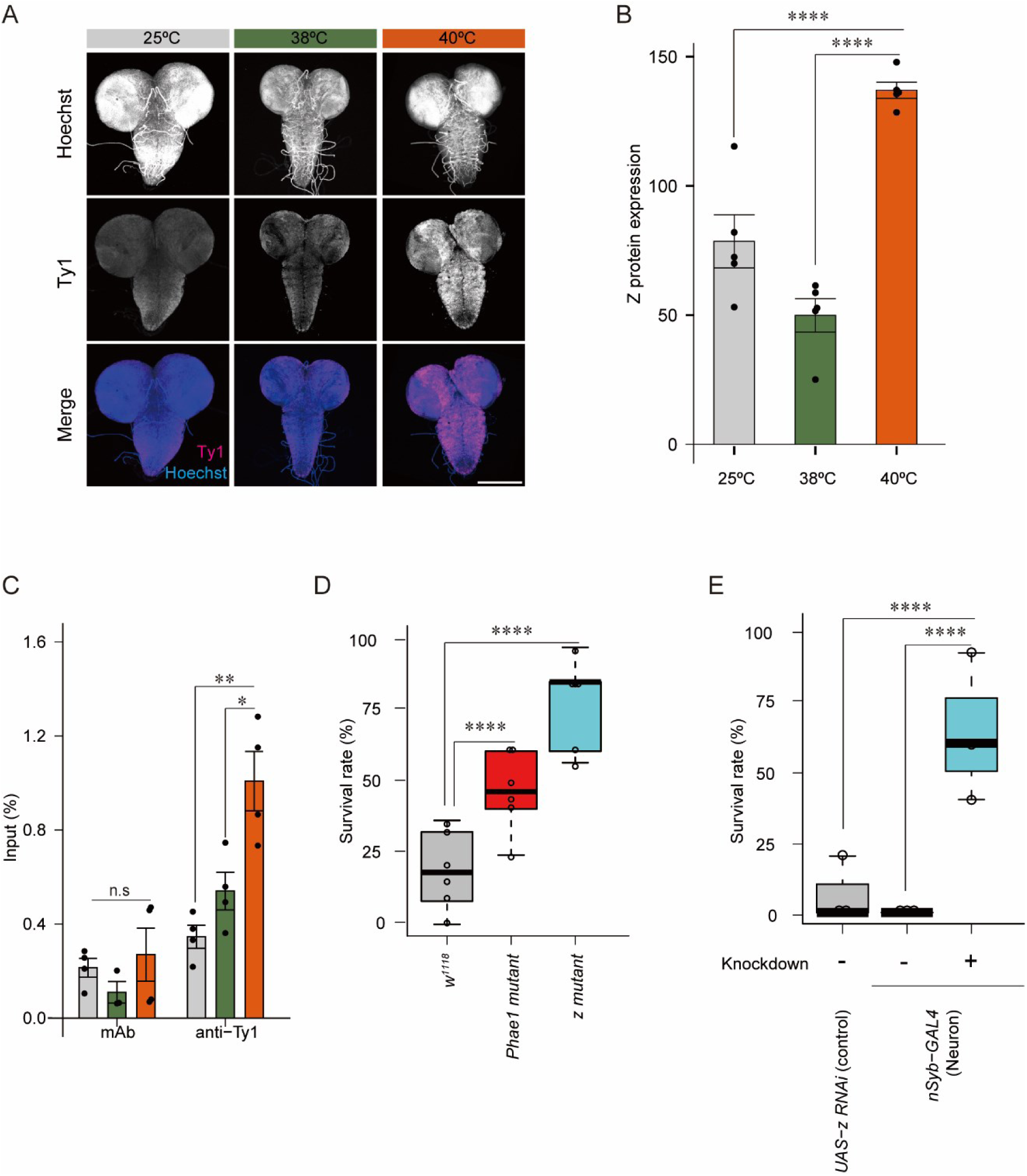
Z regulates larval survival and *Phae1* induction following heat stress exposure. (A) CNS stained with anti-Ty1 tag antibody to detect endogenous Z protein. Scale bar: 200 μm. (B) Quantification of Z protein level. \*\*\*\**P* < 0.001 (one-way ANOVA followed by Tukey’s HSD). N = 5 independent technical replicates. (C) ChIP assay shows the relative precipitation of the *Phae1* enhancer region in larval CNS following exposure to lethal heat stress. \*\*\*\**P* < 0.001 (one-way ANOVA followed by Tukey’s HSD). N = 4 independent technical replicates. (D) Survival of control (*w^1118^*), *Phae1* mutant, and *z* mutant larvae following a 30-min exposure to lethal heat stress at 40 °C. \*\*\**P* < 0.001 (Fisher’s exact test). All values are means ± SE. N = 6 independent technical replicates, n = 25 independent biological replicates. (E) Survival of larvae with or without neuro-specific *z* knockdown following exposure to a 30-min lethal heat stress at 40 °C. \*\*\*\**P* < 0.001 (Fisher’s exact test). All values are means ± SE. N = 3 independent technical replicates, n = 25 independent biological replicates.

### The neuronal mTOR-Z-Phae1 axis regulates heat stress-induced organismal death

To further investigate the upstream signaling pathways that regulate *Phae1* expression, we performed a small molecular inhibitor screen (34) in S2 cells using a luciferase reporter (*Phae1>Luc*) assay (Fig. 6A). We compared the effect of 96 different small molecule inhibitors (10 nM) on *Phae1* transcriptional activity and found that rapamycin, the well-known inhibitor of mechanistic target of rapamycin (mTOR), produced the most significant reduction in *Phae1>Luc* activity (Fig. 6B, Appendix Dataset 03). Rapamycin produced a dose-dependent suppression of *Phae1>Luc* activity (Fig. 6C), while also reducing TUNEL^+^ cell death and caspase-3 activation in S2 cells following a 30-minute exposure to heat stress at 42 °C (Fig. 6D and E). These results suggest rapamycin suppresses heat stress-induced cell death by reducing *Phae1* expression.

**Figure 6.**
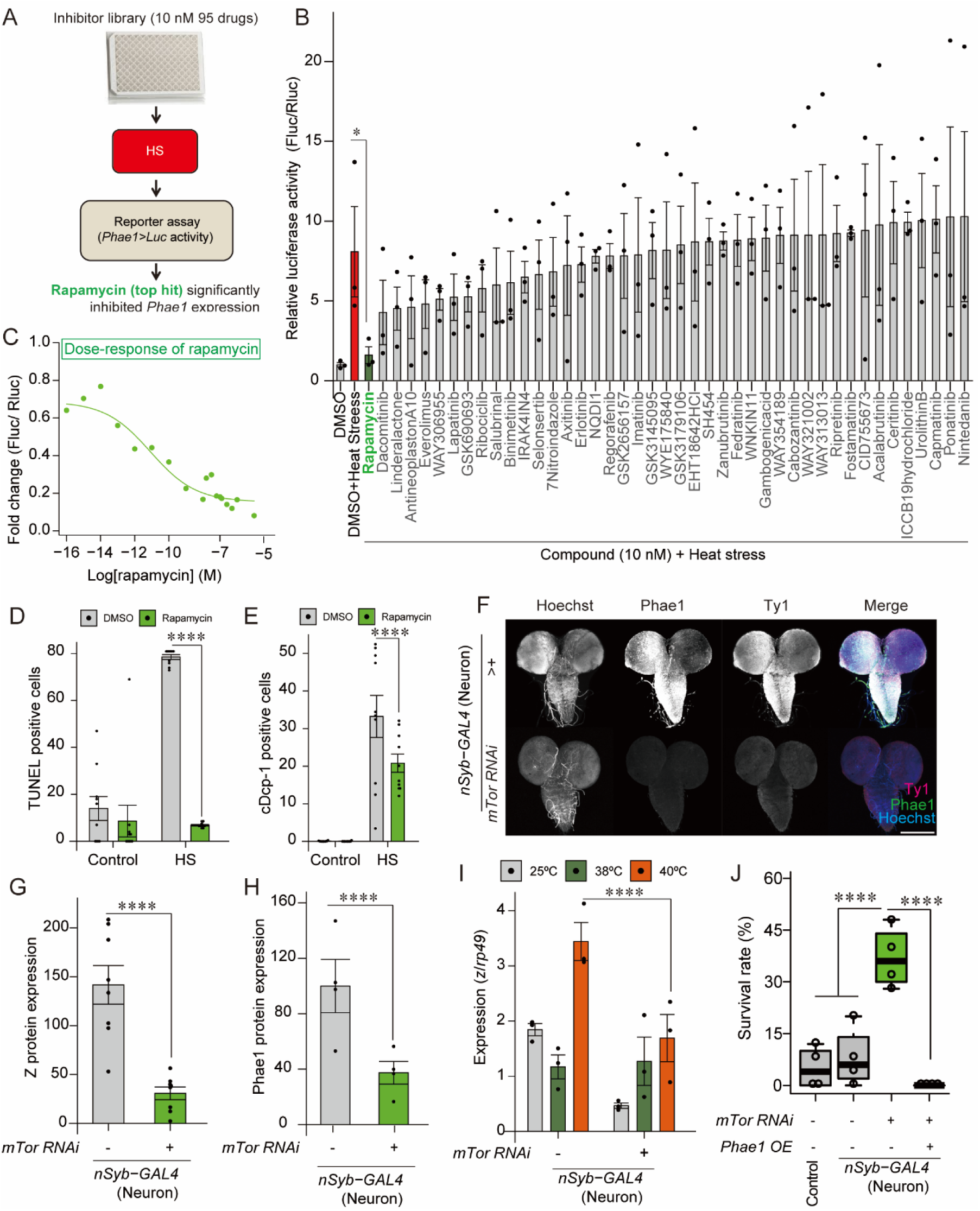
The mTOR-Z-Phae1 axis genetically regulates heat stress-induced organismal death. (A-B) Chemical screen to identify signaling pathways upstream of *Phae1* expression following heat stress exposure at 42 °C for 30-min. \**P* < 0.05 (Mann-Whitney-Wilcoxon test with Bonferroni correction). N = 3 independent technical replicates. (C) Dose-response curve for the effect of rapamycin on *Phae1* expression. (D) The effect of rapamycin on the induction of cell death following a 30-min exposure to 42 °C. \*\*\*\**P* < 0.001 (two-tailed Student’s *t* test). (E) The effect of rapamycin on caspase activation induced by a 30-minute exposure to 42 °C. \*\*\*\**P* < 0.001 (two-tailed Student’s *t* test). (F) CNS stained with anti-Ty1 antibody, anti-Phae1 antibody, and Hoechst stain for detecting Z protein in the presence or absence of neuron-specific *mTor* knockdown. Scale bar: 200 μm. (G) A quantification of Z protein level in the presence or absence of neuron-specific *mTOR* knockdown. \*\*\*\**P* < 0.001 (two-tailed Student’s *t* test). N = 5 independent technical replicates. (H) A quantification of Phae1 protein level in the presence or absence of neuron-specific *mTor* knockdown. \*\*\*\**P* < 0.001 (two-tailed Student’s *t* test). N = 5 independent technical replicates. (I) *z* gene expression in the presence or absence of neuron-specific *mTor* knockdown. \*\*\*\**P* < 0.001 (two-tailed Student’s *t* test). N = 3 independent technical replicates. (J) Survival of larvae in the presence or absence of neuron-specific *mTor* knockdown following exposure to lethal heat stress. \*\*\*\**P* < 0.001 (Fisher’s exact test). All values are means ± SE. N = 4 independent technical replicates, n = 25 independent biological replicates.

We next asked whether mTOR affects heat stress-induced organismal death by altering *Phae1* expression. We found neuron-specific knockdown of *mTor* reduced Z and Phae1 protein levels in the larval CNS following exposure to lethal heat stress (Fig. 6F), indicating that mTOR signaling induces *Phae1* expression by regulating Z levels (Fig. 6G and H). CNS-specific *mTor* knockdown also suppressed *z* expression (Fig. 6I) and increased larval survival following lethal heat stress (Fig. 6J), but this increased survival was completely abolished by neuron-specific overexpression of *Phae1*. We therefore conclude that the mTOR-Z-Phae1 axis regulates heat stress-induced organismal death.

## Discussion

This study identifies *Phae1* as a key mediator promoting heat stress-induced organismal death. *Phae1* was initially identified as a gene located near *Drab6*, which encodes a Ras-like small GTPase involved in vesicle trafficking (23). Coincidentally, *Phaedra* was named after a woman in Greek mythology who committed suicide (35).

*Phae1* encodes a protein homologous to members of the kallikrein family (23) of secreted serine proteases, but its biological function in insects has not yet been otherwise reported. The *Phae1* gene is evolutionally conserved in Diptera and Hymenoptera according to the MEROPS database (https://www.ebi.ac.uk/merops/cgi-bin/sequence_features?mid=S01.B64) (36). Our experiment using *Phae1* transgenes with three alanine substitutions in the putative Phae1 active site amino acid residues (H76A:D125A:S223A) indicated that Phae1 serine protease activity is critical for heat stress-induced organismal death. In mammals, some proteases of the kallikrein family are involved in the induction of autophagy and apoptosis (37). Since Phae1 is required for heat stress-induced neuronal cell death and caspase activation, it is likely that Phae1 shares similar functional properties with mammalian kallikrein proteases. In addition, Kallikrein-related peptidase-7 cleaves Caspase-14 and regulates its maturation (38). We therefore hope to examine in a future study whether any of the *D. melanogaster* caspases are substrates of Phae1.

Our data suggested that the trihelix transcription factor Z binds the *Phae1* enhancer, inducing *Phae1* expression to regulate neuronal cell death following exposure to lethal heat stress. Z was originally identified as an eye color-related gene, because *z* mutant flies have brown eyes (39). Another previous study showed that Z regulates cell death induction in *D. melanogaster* salivary glands during pupariation, along with Ecdysone signaling (40). It is therefore possible that Z is a key regulator of cell death induction in various tissues.

We also found that *z* expression is regulated by mTOR, which acts as a nutrient sensor (41, 42). Starvation stress reduced *Phae1* expression probably by suppressing mTOR (SI Appendix Fig. S2B). Neuronal mTOR regulates cell death induction in *D. melanogaster* (43), and mTOR inhibition suppresses a form of stress-dependent, apoptosis-like cell death along with caspase activation in mammalian cells (44, 45). Consistent with these previous studies, our findings show that rapamycin and neuron-specific *mTor* knockdown reduce caspase activation, inhibit cell death, and increase larval survival following lethal heat stress. Together, our findings suggest that the genetically regulated mTOR-Z-Phae1 axis can determine whether an animal survives or succumbs to heat stress-induced death.

Recent studies have proposed that stress-induced organismal death is genetically regulated (46, 47). In this study, we found that lethal heat stress activates the expression of proteases and chitin metabolism-related genes, which are associated with tissue degeneration in dying insects (48, 49). Thus, our results support the hypothesis that a specific genetic mechanism promotes heat stress-induced organismal death. To individuals in a group, stress-induced organismal death can offer the benefit of eliminating potentially dangerous individuals, such as those with infections, and reducing competition for space and food (50). In some organisms, such as the budding yeast *Saccharomyces cerevisiae* and the nematode *Caenorhabditis elegans,* such an “altruistic” sacrifice can increase survival under harsh conditions (51, 52). The hypothesis that individual organismal death can contribute to a thriving population has not been completely proven in most animals, but mathematical simulations have suggested that stress-induced organismal death can be adaptative when the cost of cohort competition/danger is sufficiently large (53). In this respect, it will be interesting to examine whether and how the mTOR-Z-Phae1 axis contributes to *D. melanogaster* population fitness.

It is noteworthy that genes like *D. melanogaster Phae1*, *z*, and *mTor* are present in mammalian genomes. As mentioned above, Phae1 shows homology with mammalian kallikrein. The helix-turn-helix DNA binding domain of Z is also found in the mammalian nuclear apoptosis-inducing factor (NAIF) (54). Moreover, mTOR signaling is evolutionarily conserved in various animals, including mammals (55), and its inhibition can increase lifespan and stress resistance across several species (56, 57). Therefore, it will be interesting to determine just how widely the mTOR-Z-Phae1 axis is conserved.

## Materials and Methods

This study included RNA-seq, qRT-PCR, gel shift assays, multiple staining methods, and luciferase reporter assays. Detailed descriptions of the materials and methods used in this study are provided in the Supporting Information (SI). All primers are listed in Table S1, and all small molecule inhibitors appear in Table S2.

## Supporting information

Spplement-AI

## Author Contributions

T.M & R.N wrote manuscript, T.M performed experiments, M.R, supported experiments, S.K & A.N constructed transgenic lines, T.M, H.M, T.K, Y.H, R.N designed experiments.

## Acknowledgments

We thank the following for their advice and generosity with their resources: N. Okamoto, K. Iwasaki, K. Takahashi, T. Yoshiga, M. Tokuda, and T. Tujita. We thank the Bloomington Stock Center (NIH P40OD018537), the Kyoto Stock Center, the Vienna Drosophila RNAi Center, and the Developmental Studies Hybridoma Bank (created by the NICHD of the NIH and maintained at The University of Iowa, Department of Biology, Iowa City, IA 52242) for fly stocks and antibodies. This work was supported by JSPS KAKENHI (19J12272, 22KJ0339, 24K18156) grants to T.M., by a Promotion of Development of a Joint Usage/ Research System Project: Coalition of Universities for Research Excellence Program (CURE) from the MEXT (JPMXP1323015486) grant to A.N., and by support from the program of Joint Usage/Research Center for Developmental Medicine, Institute of Molecular Embryology and Genetics, Kumamoto University (K24-15) to R.N.. T.M. was also a recipient of a JSPS Young Investigator fellowship.

