## Supplementary material for "Stress-induced organismal death is genetically regulated by the mTOR-Zeste-Phae1 axis": Spplement-AI

[a]Life Science Center for Survival Dynamics, TARA, University of Tsukuba, Japan, [b]Analytical Research Center for Experimental Sciences, Saga University, Japan, [c]Department of Applied Biological Science, Saga University, Japan, [d]Department of Biological Science and Technology, Tokyo University of Science, Japan, [e]Invertebrate Genetics Laboratory, National Institute of Genetics, Japan, [f]Institute of Molecular Embryology and Genetics, Kumamoto University, Japan

*To whom correspondence may be addressed.

**Supporting Information Text**

**Materials and Methods**

**Animals and heat stress.** Fly stocks were maintained on a standard diet (0.55 g agar, 10.0 g glucose, 9.0 g cornmeal, 4.0 g yeast extract, 300 µl propionic acid, 350 µl 10 % butylparaben in 70% ethanol, and 100 ml water). All experiments were conducted at 25 °C under a 12:12-hour light/dark cycle (1). Third instar larvae were used for all experiments. Test larvae were exposed to heat stress by placing individuals in a glass vial (55 mm × 22 mm i.d.) that was then dipped in a water bath (Taitec, #SDM-B) held at 37–40 °C. All experimental heat treatments lasted 30 min. Controls were exposed to 25 °C. For the chemical screen, *Drosophila* larvae were reared for 12 hours on foods containing each chemical. Larval survival rates were determined by observing the activity of the larval dorsal vasculature of 25 biologically independent larvae.

**Establishment of transgenic flies.** For the construction of the UAS vectors, the *Phae1* coding sequence (CDS) was obtained via PCR from larval *D. melanogaster* central nervous system (CNS)-derived complementary DNAs (cDNAs) and primers (Eco RI_*Phae1* Fw and Xba I_*Phae1*_Rv; see Table S1). The PCR product and pUAST plasmids (DGRC, #1000) were digested with *Eco* RI and *Xba* I. The digested PCR product and pUAST plasmids were then ligated with Ligation High Ver.2 (TOYOBO, #LGK-201), producing the final *UAS-Phae1* vector. The *Phae1-T2A-GAL4* strain was established by WellGenetics Co. Ltd. (https://wellgenetics.com).

*Phae1* knock-in mutant animals were established by introducing mutations and tags to the *Phae1* CDS using the CRISPR-Cas9 method as previously described (2). The gRNA target sites were determined using the gRNA target prediction tool Fly Cas9 (https://shigen.nig.ac.jp/fly/nigfly/cas9/cas9TargetFinder.jsp). The following pairs of oligonucleotides were used (Table S1): *Bbs I_Phae1 gRNA_F1* and *Bbs I_Phae1 gRNA_R1* for $Phae1^{WT}$*::HA*, *Bbs I_Phae1 gRNA_F2* and *Bbs I_Phae1 gRNA_R2* for $Phae1^{EI\ mutant}$*::HA*. The pDCC6 plasmid (Addgene, #59985) was used to construct the vectors for expressing the gRNA and Cas9. A single guide RNA (sgRNA) targeting the Phae1 protein coding site (CDS) was introduced to replace the Phae1 CDS, resulting in a HA-tagged Phae1 protein with or without mutations in the Phae1 active site ($Phae1^{WT}$*::HA* or $Phae1^{EI\ mutant}$*::HA*).

A knock-in fly strain expressing Ty1-tagged Z was established by editing the endogenous *z* gene using the CRISPR-Cas9 method with *Bbs I_Z gRNA_F* and *Bbs I_Z gRNA_R*. The pTWIST plasmids, including the homology arms corresponding to parts of genomic sequences surrounding the *z* locus (SI Appendix, Dataset S03), were obtained from Twist Bioscience (https://twistbioscience.yokohama/). All transgenic strains were established via injection of vector plasmids into embryos.

**RNA isolation.** Total RNA was extracted from *Drosophila* tissues using TRIzol (Invitrogen, #15596026) according to the manufacturer's protocol. Samples contained tissues dissected from 20 individuals for the larval CNS and wing discs or 10 individuals for other tissues. Total RNA quality was checked by NanoDrop2000c (Thermo Scientific). Purified RNA was used for RNA-sequencing and qRT-PCR analysis.

**RNA-sequencing analysis.** RNA-seq was performed to analyze gene expression in *D. melanogaster* larval fat bodies following exposure to non-lethal heat stress (38 °C) or lethal heat stress (40 °C). Libraries were sequenced on an Illumina Hiseq4000 sequencer by Genewiz (www.genewiz.com). The concentrations of the prepared library solutions were measured by Qubit3.0 Fluorometer and dsDNAHS AssayKit (ThermoFisher Scientific). Library quality was checked by Bioanalyzer and HighSensitivityDNAKit (Agilent Technologies). Reads were aligned to the *Drosophila melanogaster* genome (Genome assembly, #BDGP6, https://useast.ensembl.org/Drosophila_melanogaster/Info/Index) using TopHat (version 2.1.2). Raw reads are accessible at the DNA Data Bank of Japan (DDBJ) (accession number; DRR620746-DRR620751 (x18976002-0001_Run_0001-0006)).

**qRT-PCR.** cDNA was synthesized with the ReverTra Ace qPCR RT kit (Toyobo, # FSQ-101). qPCR analysis of each gene was carried out using a CFX-Duet instrument (BIO-RAD) through 50 cycles of 95 °C for 10 s, 55 °C for 10 s, and 72 °C for 20 s. PCR specificity was confirmed by sequencing the PCR products and performing a melting curve analysis for each data point. Data were normalized to the expression of *rp49* determined in duplicate by reference to a serial dilution calibration curve (3). The primer pairs used for PCR in this study are listed in Table S2.

**Electrophoretic mobility shift assay (EMSA).** A GST-His tagged polypeptide of 76 amino acids, corresponding to the Z DNA binding domain and its surrounding regions (ZDBD, 37.8kDa)(4) was obtained from an *E.coli* BL21 strain (Agilent, # 200132) with the pCold-GST system (TaKaRa, #3372) and then purified using AKTA-Go (Cytiva). The quality of the ZDBD protein was verified via SimpleWestern blotting (Protein simple, Abby; # 004-680-PS-T006, Appendix Fig. S7). The 560 bp *Phae1* enhancer region was used as a free DNA probe. Transient mutations were introduced into the Z-binding motif of the *Phae*1 enhancer region using the KOD -Plus- Mutagenesis Kit (TOYOBO, #SMK-101) and the resulting sequence was used as the mutated DNA probe. Anti-His tag antibody (1:50, MBL, #M089-3) was used for a super shift assay. 0–5 μM samples of ZDBD were incubated with 5 ng/samples of *Phae1* enhancer region DNA probe with or without transient Z-binding motif mutations in reaction buffer (10 mM Tris-HCl, 1 mM DTT, 50 mM NaCl) at 25 °C for 30 min (5). 8% Native-PAGE gels were prepared (30 w/v% acrylamide/bis mixed solution (FUJIFILM, #018-25625); 2.7 mL 5x TBE (0.445 mol/l Tris-borate, 10 mmol/l EDTA); 1 mL, DW; 6.2 mL, 10 w/v% ammonium persulfate (Kanto Chemical, #01307-00); 0.1 mL, N,N,N',N'-tetramethylethylenediamine (Fujifilm Wako, #202-04003); 10 μL). Samples were applied and subjected to TBE-PAGE (125 V const, 50 min). Free and shifted DNA bands were detected with MIDORI Green Xtra (1:10,000, NIPPON genetics, #MG10).

**Immunohistochemistry.** Tissues were dissected from test larvae and fixed in 4% paraformaldehyde (PFA) in PBS for 30 min at room temperature. After washing in PBS containing 0.1% Triton-X (PBT) for 30 min, the tissue samples were blocked in PBS containing 2% bovine serum albumin (BSA) for 1 hour at room temperature and then incubated with primary antibodies at 4 °C overnight. Rabbit anti-Phae1 IgG (this study, 1:500), mouse anti-Repo IgG (DSHB, #8D12, 1:200), rabbit anti cleaved Dcp-1 (Cell signaling, #9578, 1:1,000), and mouse anti-Ty1 IgG (Sigma, #SAB4800032, 1:10,000) primary antibodies were used. Hoechst33342 (Sigma, #2261, 10 μM) was used to visualize nuclei. After three 30-min washes in PBT, the samples were incubated with Alexa Fluor 488 or 546-conjugated secondary antibodies (Thermo Fisher Scientific, #R37116; # R37117; #R37121, 1:500) for 1 hour.

**TUNEL labeling and cell counting.** The *In Situ* Cell Death Detection Kit with TMR red (Roche, #12156792910) was used for TUNEL labeling. Fixed CNS samples were subjected to three 30-min washes in PBT. The tissue samples were added to 25 μL labeling solution and incubated at 37 °C for 1 hour while being protected from light. After three 10-min washes in PBT, the samples were mounted in FluorSave reagent (Merck Millipore, #34578). The samples were then visualized using a Zeiss LSM 900 confocal microscope as previously described (1). The number of TUNEL-positive cells and cleaved Dcp-1 positive cells were counted in confocal microscopy images with Cellpose2.0 (6).

**ChIP assay.** The *Drosophila* ChIP-seq kit (Chromatrap, # 500274) was used according to the manufacturer's protocol to verify the interaction between Z and the *Phae1* enhancer region. Twenty-five *D. melanogaster* larval CNS were used for each sample. The quality of the genomic DNA was checked by NanoDrop2000c (Thermo Scientific).

**Protein 3D structure.** AlphaFold 3 (https://alphafoldserver.com/about) was used for the prediction of protein 3D structure (7). UCSF Chimera X (https://www.cgl.ucsf.edu/chimerax/) was used for editing protein 3D structures (8). Based on the 3D structures predicted by AlphaFold 3, ortholog candidates were identified using the protein ortholog prediction tool Foldseek (https://search.foldseek.com/search) and the MEROPS Peptidase Database (https://www.ebi.ac.uk/merops/cgi-bin/sequence_features?mid=S01.B64) (11).

**Luciferase reporter assays and chemical screening.** The dual-Luciferase Reporter Assay System (Promega, #E1910) was used to identify stress response elements in the *Phae1* promoter as previously described (9). The inserted enhancer regions were prepared by PCR using the primers *Kpn I_Luc_Fw 530 bp, Kpn I_Luc_Fw 540 bp, Kpn I_Luc_Fw 550 bp*, and *Xho I_Luc_Rv*, which appear in Table S1. The pGL4.23 (Promega, #9PIE841) plasmid, containing the *Phae1* enhancer region, was transfected into S2 cells. pGL4.70 (Promega, #E6881) was used as a control vector for the dual luciferase reporter assays. Luciferase activity was measured 72 hours after transfection. For the chemical screen, a small molecular inhibitor library was obtained from Sellechchem (https://www.selleckchem.com/). *Phae1>Luc* cells were treated with 10 μM of each compound for 8 hours, and then luciferase activity was measured. All chemicals used for the chemical screen are listed in SI Appendix, Dataset 02.

**Measurements of GFP fluorescence intensity.** For the measurements of GFP fluorescence signals, ten *Phae1>GFP* larvae were collected and washed with distilled water. The larvae were then homogenized in 200 µL lysis buffer (100 mM Tris, pH 8.0, 100 mM borate, 0.1 M EDTA, 1% Triton-X) at 4 °C. The homogenates were centrifuged at 12,000 × *g* for 5 min at 4 °C, and the supernatants were used as samples to measure the relative intensity of GFP fluorescence. Relative GFP fluorescence intensities were measured using a Multimode Detector DTX880 (BeckmanCoulter) at Ex/Em=488/595 nm. Protein quantification was carried out using the Bradford method with BSA as a standard (10).

**Statistical analysis.** Statistical analyses were carried out using two-tailed Student's *t* tests, one-way ANOVA followed by Tukey's multiple comparison tests, Fisher's exact tests, and Mann-Whitney-Wilcoxon test with Bonferroni correction. *P*-values of less than 0.05 were considered statistically significant. All statistical analyses were performed using JMP 9.0.2 (SAS Institute). All data used for the statistical analyses are available in SI Appendix, Datasets S01-S03.

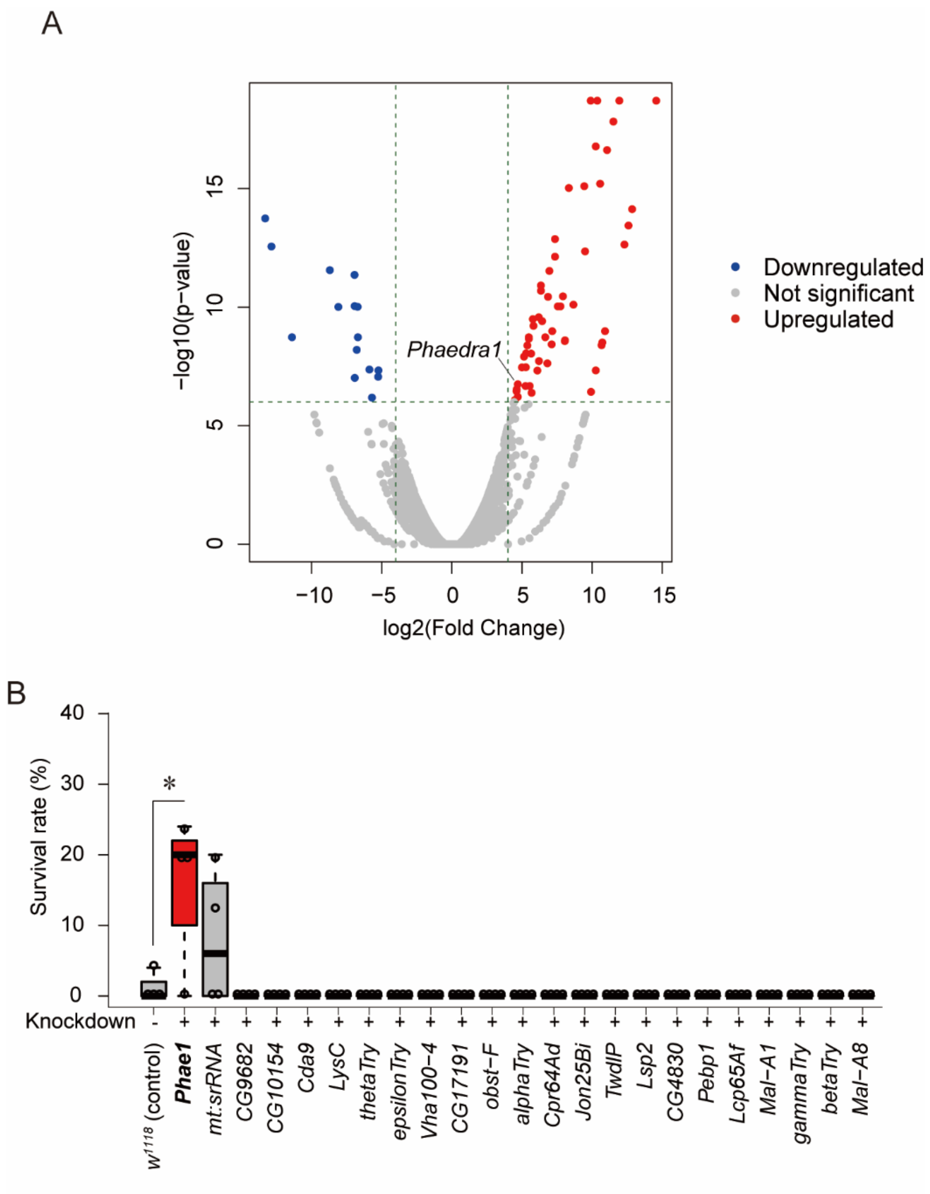


**Fig. S1.** Identifying death mediator gene candidates. (A) Volcano plot of the RNA-seq data for identifying death mediator gene candidates. The red circles indicate upregulated genes (fold change > 2, FDR < 0.000001), the blue circles indicate down-regulated genes (fold change < 1/2, FDR < 0.000001), and the gray circles indicate genes unchanged following lethal heat stress at 40 °C. (B) Survival of larvae with knockdown of Phae1 or other lethal stress-induced genes after a 30-min exposure to lethal heat stress at 40 °C. All values are means ± SE. **P* < 0.05 (Fisher's exact test). All values are means ± SE. *****P* < 0.001 (Fisher's exact test). N = 25 independent biological replicates, n = 4 independent technical replicates.

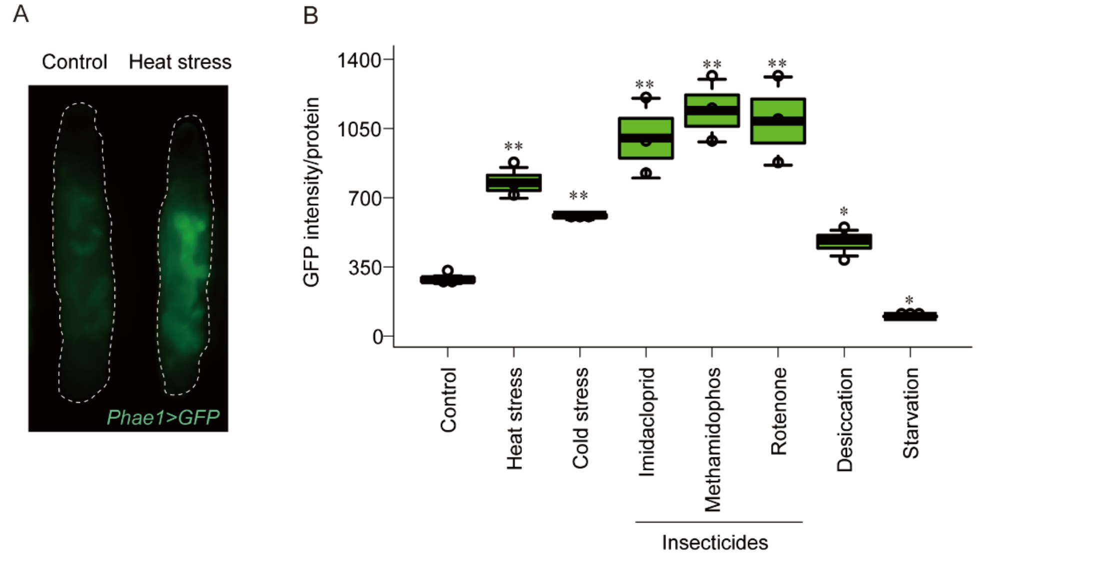


**Fig. S2.** *Phae1* expression is upregulated by various stressors. (A) Fluorescence microscopy image of the transgenic *Phae1>GFP* reporter strain with or without a 30-min exposure to lethal heat stress at 40 °C. Dotted lines encircle the larval bodies. (B) Quantification of Phae1>GFP intensity from larvae exposed or unexposed to the indicated stressors. Control; unstressed larvae, Heat stress; 40 °C for 30 min, Cold stress; 4 °C for 12 h, Imidacloprid; 100 μM for 12 h, Methamidophos; 100 μM for 12 h, Rotenone; 100 μM for 12 h, Desiccation; relative humidity 15% for 5 h, Starvation; 2% agar for 12 h. All values are means ± SE. $^{**}P < 0.001$, $^{*}P < 0.05$ (two-tailed Student's t test) compared with unstressed control larvae at 25 °C. N = 10 independent biological replicates, n = 3 independent technical replicates.

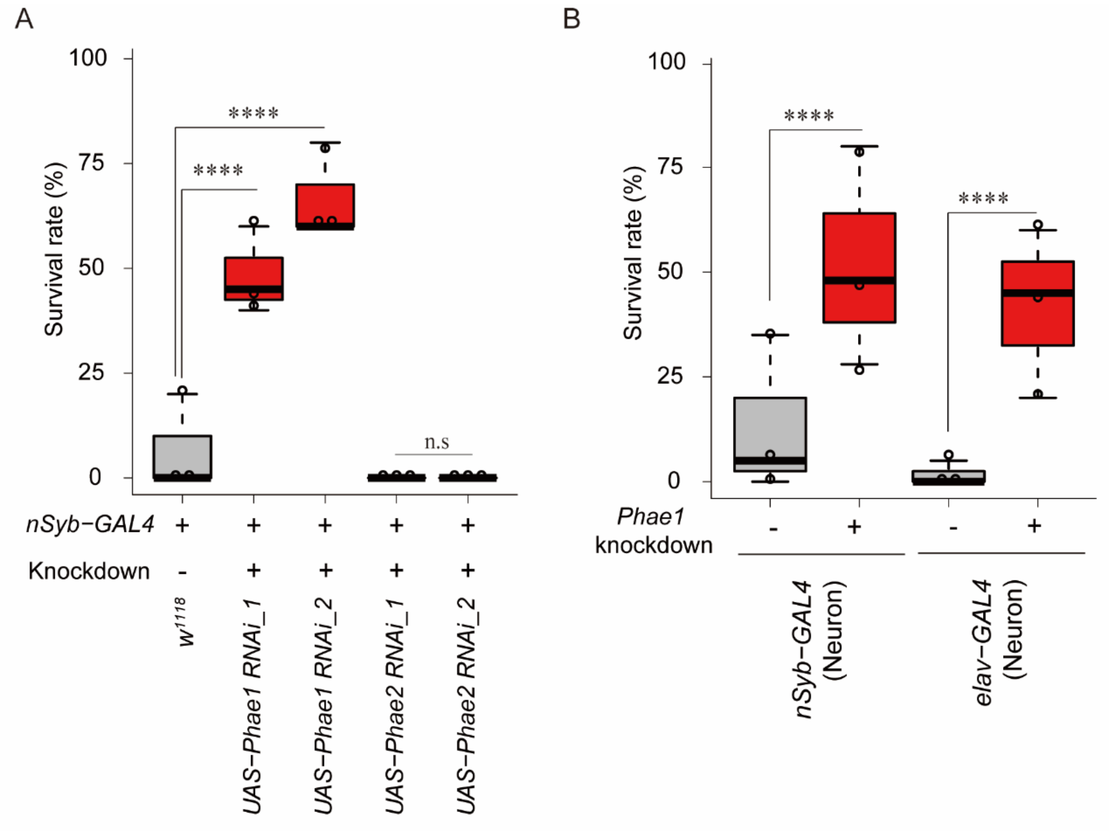


**Fig. S3.** Survival of animals with neuron-specific knockdown of *Phae1* or *Phae2*. (A) Survival of control (*nSyb-GAL4*>+) and *Phae1* or *Phae2* knockdown larvae exposed to lethal heat stress. All values are means ± SE. **P* < 0.05 (Fisher's exact test). N = 25 independent biological replicates, n = 3 independent technical replicates. (B) Survival of larvae with or without pan-neuronal *Phae1* knockdown. N = 25 independent biological replicates, n = 3 independent technical replicates. All values are means ± SE. **P* < 0.05 (Fisher's exact test).

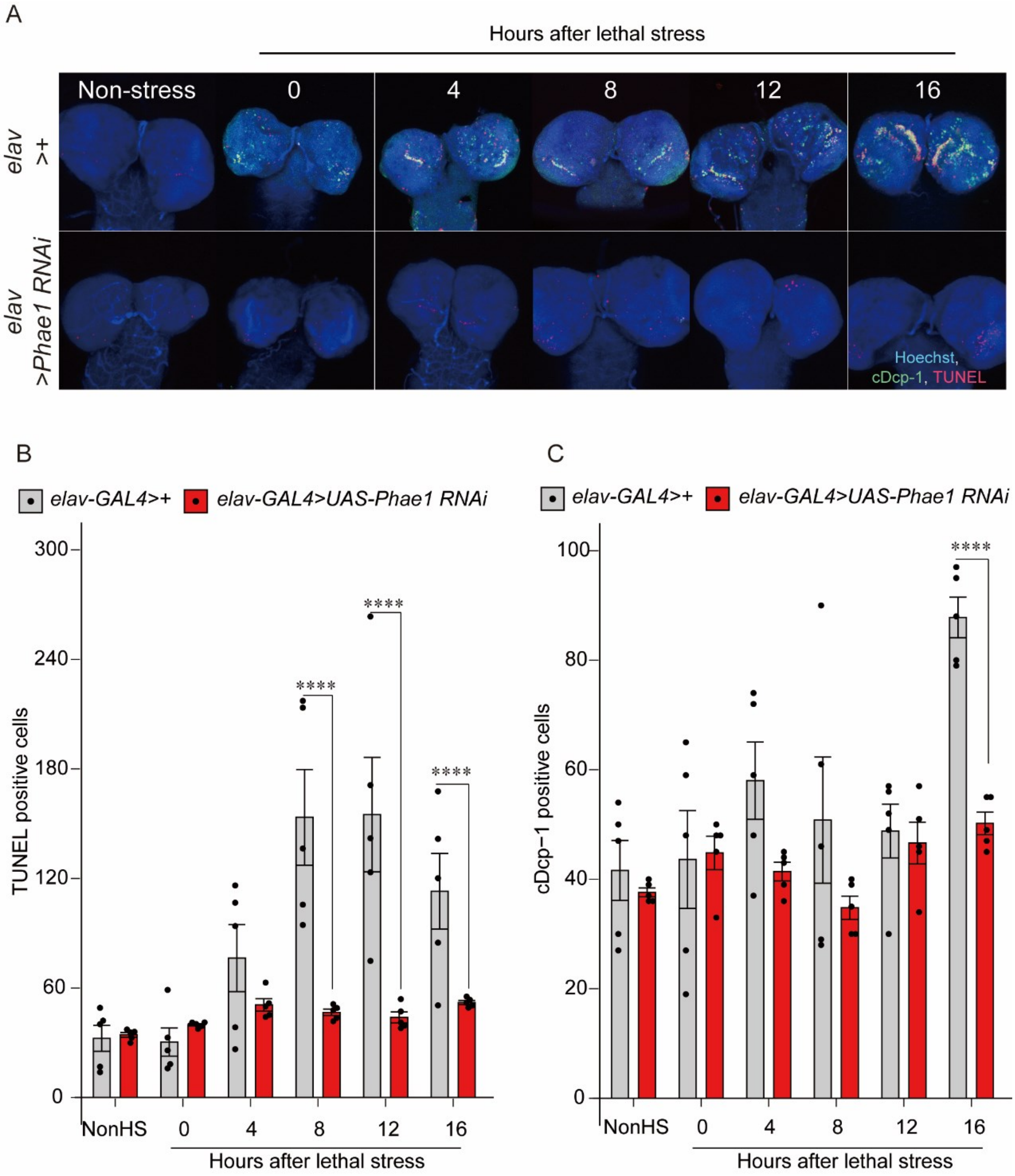


**Fig. S4.** Neuronal *Phae1* knockdown reduces caspase activation and cell death. (A) Confocal images of the CNS stained with Hoechst (nuclear DNA), anti-cleaved Dcp-1 (activated caspase-3), and TUNEL (dead cells). (B, C) Quantification of TUNEL-positive cells (B) and cDcp-1-positive cells (C) in the CNS of control and neuronal *Phae1* knockdown larvae. ****$P$ < 0.001 (two-tailed Student's t test). N = 10 independent biological replicates, n = 5 independent technical replicates.

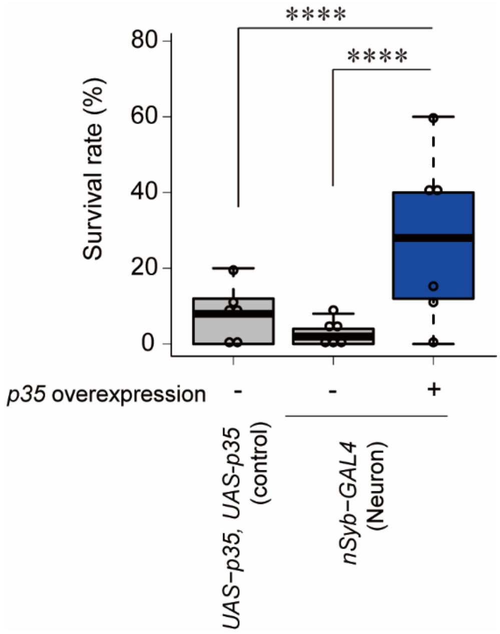


**Fig. S5.** Neuronal caspase inhibition increases survival following exposure to lethal heat stress. All values are means ± SE. ****$P < 0.001$ (Fisher's exact test). N = 25 independent biological replicates, n = 6 independent technical replicates.

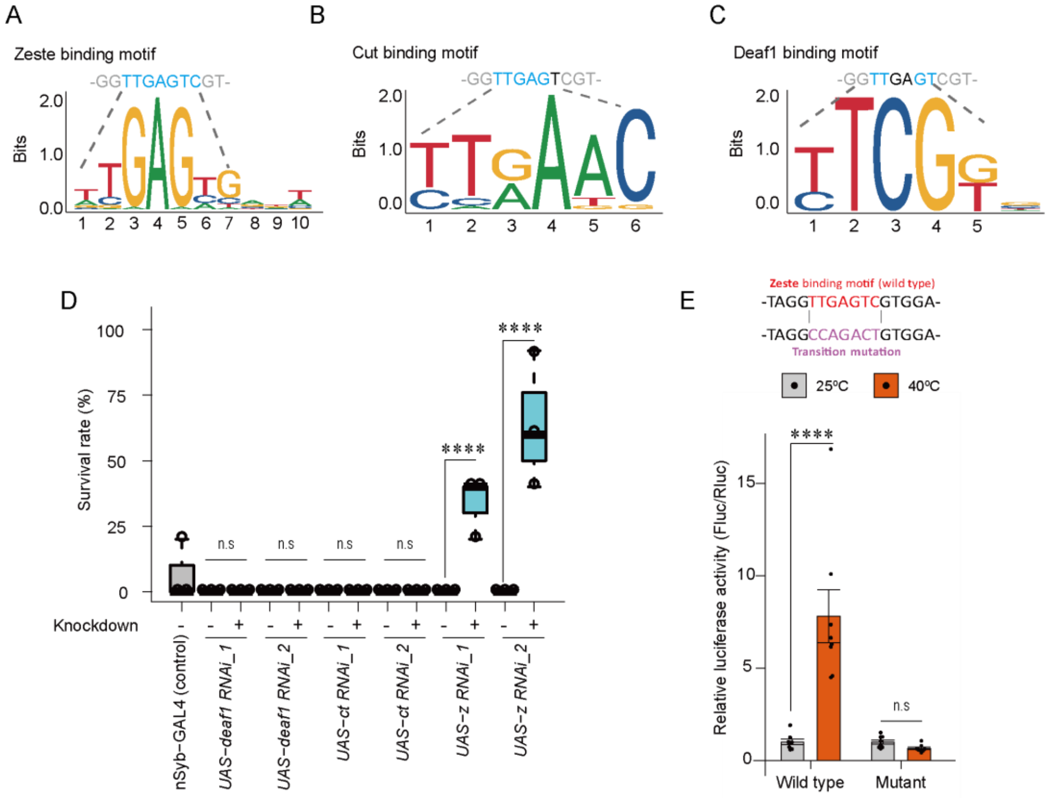


**Fig. S6.** Z, but not Cut or Deaf1, affects both Phae1 expression and survival following heat stress. (A) Z binding motif. (B) Cut binding motif. (C) Deaf-1 binding motif. Transcription factor binding motifs LOGOs were generated by JASPAR. (D) Lethal heat stress survival of control larvae (*nSyb-GAL4*) vs. larvae with neuron-specific knockdown of *z*, *cut*, or *deaf1*. All values are means ± SE. ****$P < 0.001$ (Fisher's exact test). N = 25 independent biological replicates, n = 3 independent technical replicates. (E) Luciferase activity in S2 cells transfected with wild-type *Phae1* enhancer reporter or a modified enhancer reporter containing a mutated Z-binding motif. ****$P < 0.001$ (two-tailed Student's t test). N = 5 independent technical replicates.

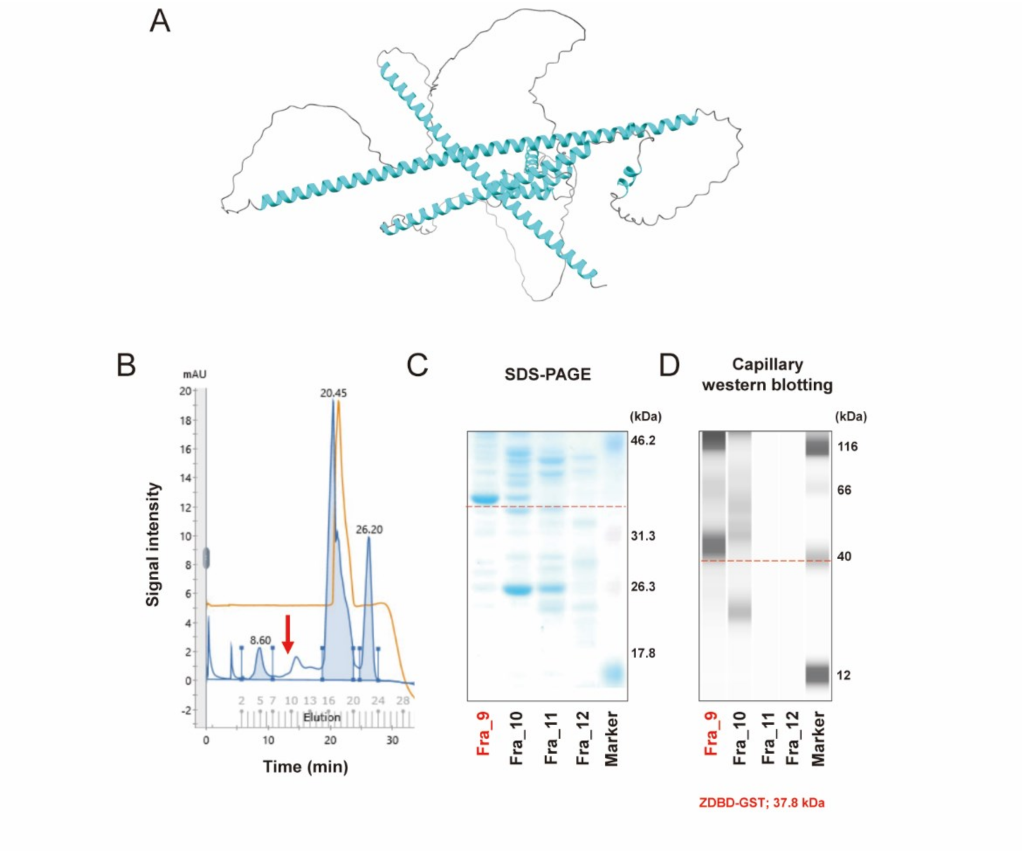


**Fig. S7.** Preparation of a recombinant Z DNA binding domain (ZDBD) conjugated to a GST tag. (A) Z protein 3D structure created by Alphafold-3 and edited by Chimera X. The blue area indicates the helix-turn-helix DNA binding domain. (B) Purification of recombinant Z protein by AKTA-Go. (C) SDS-PAGE for checking Z protein quality. (D) Simple western blotting for checking Z protein.

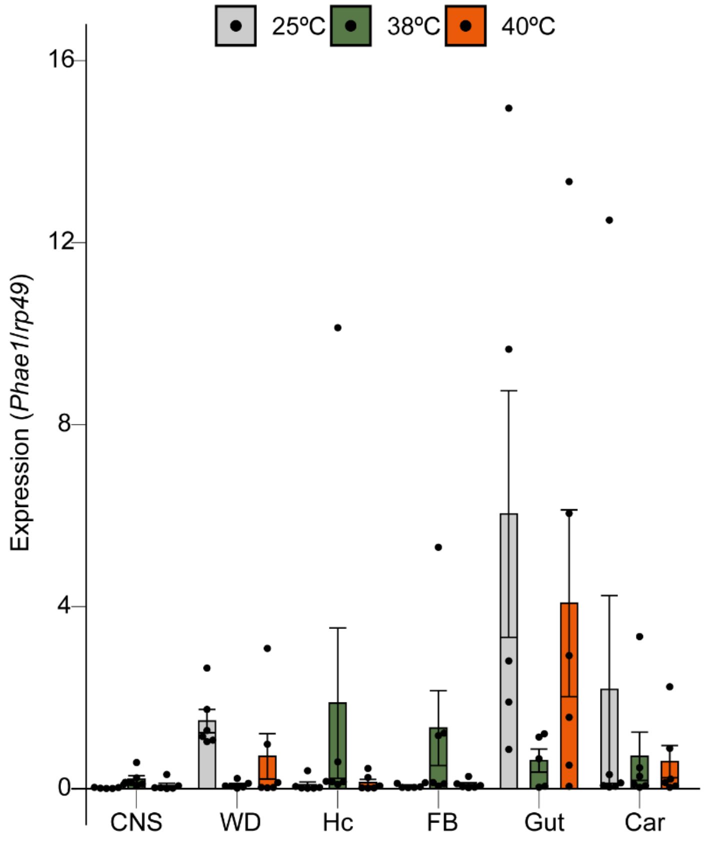


**Fig. S8.** Mutation of *z* reduces the heat stress-induced increase in *Phae1* expression in various tissues (CNS; central nervous system, WD; wing disc, Hc; hemocyte, FB; fat body, Gut, Car; carcass). The gray, green, or orange bars indicate exposure to control (25 °C), non-lethal stress at 38 °C, or lethal stress at 40 °C, respectively. N = 5 independent technical replicates.

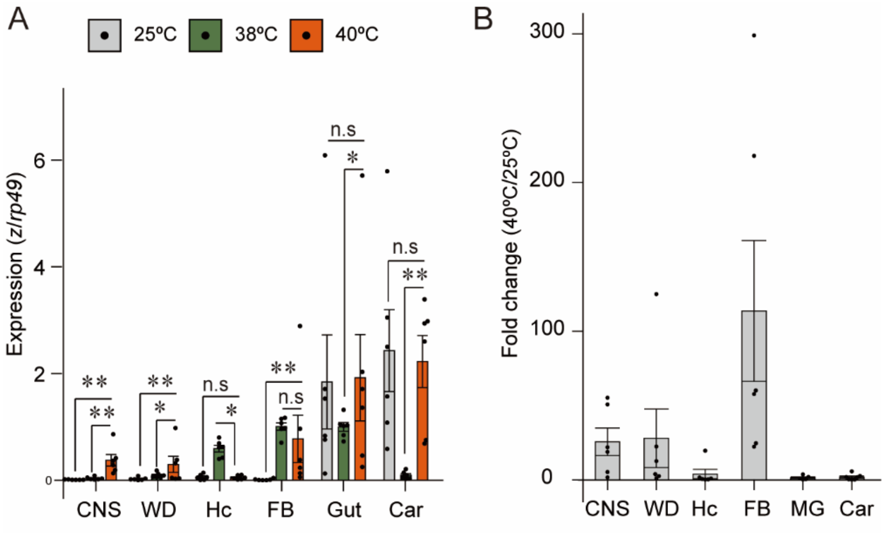


**Fig. S9.** Lethal heat stress induces *z* expression in several tissues. (A) *z* expression in various tissues exposed to the indicated temperatures. $^*P < 0.05$, $^{**}P < 0.01$ (one-way ANOVA followed by Turkey HSD). N = 6 for each sample. (B) Fold change in *z* expression for each tissue. N = 5 independent technical replicates.

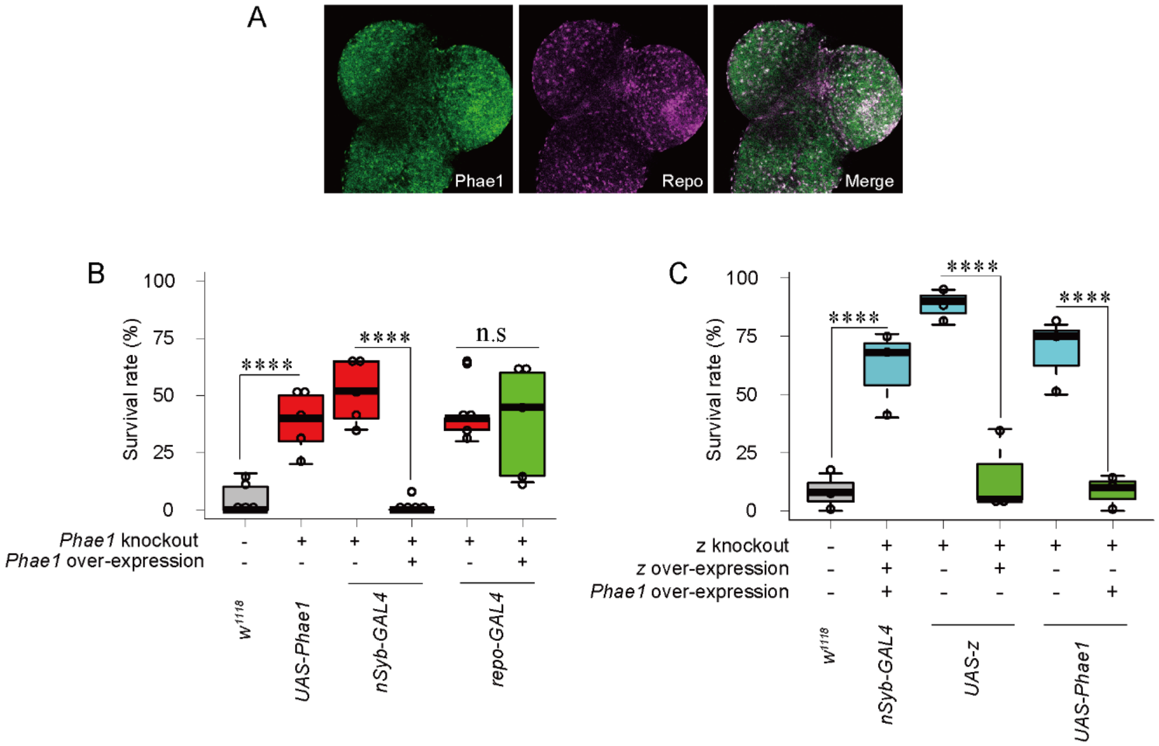


**Fig. S10.** The increased resistance to heat stress conferred by *Phae1* or *z* loss-of-function mutations is blocked by neuronal overexpression of *Phae1* or *z*. (A) Confocal images of Repo-positive cells and Phae1-positive cells in the CNS. (B) Neuronal *Phae1* overexpression in the CNS in the *Phae1* mutant background. All values are means ± SE. *****P* < 0.001 (Fisher's exact test). N = 25 independent biological replicates, n = 5 independent technical replicates. (C) Neuronal overexpression of *z* or *Phae1* in the *z* mutant background. All values are means ± SE. *****P* < 0.001 (Fisher's exact test). N = 25 independent biological replicates, n = 3 independent technical replicates.

**Table S1.** Primers and oligonucleotides used for vector construction.

| Target | Primer sequence (restriction enzyme cutting site) |
|---|---|
| *Eco RI_Phae1 Fw* | **CGGAATTCACC**GTTAGCGAACGGACAGCA |
| *Xba I_Phae1_Rv* | **CGTCTAGACTA**TTAGGACACCTGCTGGTTAGC |
| *UAS vector_Fw for genotyping* | AACCAAGTAAATCAACTGC |
| *UAS vector_Rv for genotyping* | ATCTCTGTAGGTAGTTTGTC |
| *Bbs I_Phae1::HA gRNA_F1* | **CTTC**GCGACTATTTAGGACACCTGC |
| *Bbs I_Phae1::HA gRNA_R1* | **AAAC**GCAGGTGTCCTAAATAGTCGC |
| *Bbs I_Phae1::HA gRNA_F2* | **CTTC**GATTGGTCAGGCAGTGGGCAG |
| *Bbs I_Phae1::HA gRNA_R2* | **AAAC**CTGCCCACTGCCTGACCAATC |
| *Kpn I_Luc_Fw_550bp* | **CGGGTACC**GTTGAGTCGTGGAAAATTAC |
| *Kpn I_Luc_Fw_540bp* | **CGGGTACC**GGAAAAATTACGGTCTCTGC |
| *Kpn I_Luc_Fw_530bp* | **CGGGTACC**GGTCTCTGCTTCTTCTTCTTTAA |
| *Xho I_Luc_Rv* | **GCGCTCGAG**ACCTTTATCGCGGCACGTTT |
| *Eco RI_ZDBD_Fw* | **CGGAATTCGCC**ATCCCTGTACGGATACTGTTTG |
| *Xho I_ZDBD_Rv* | **GCGCTCGAGCT**CACCCCGCGCTTCACC |
| *Bbs I_z gRNA_F* | **CTTC**GTGCTCCACTTCATGAGGTC |
| *Bbs I_z gRNA_R* | **AAAC**GACCTCATGAAGTGGAGCAC |

**Table S2.** PCR primers

| Gene Symbol | Fw | Rv |
|---|---|---|
| *CG34282* | GACATCTATGCCGAGCCCAA | CTGCTCCATGAAGGCGTAGT |
| *CG6295* | GAGACCAAGAACCGCATGGA | GCGGACACCGTAGTTGATGA |
| *CG7290* | TGTGTGACCAGGATGGAAGC | GGCGTTTACAAAGGTCGCAG |
| *CG5084* | CACCCAGAAGAACGGCGATA | GGACACAGTGCAGTTCAGGA |
| *CG17826* | ACCCTCGGGAACTTATTGCG | GCCAATCGCAGGATTTGACC |
| *CG14300* | GTACTGGCTGTGCGAGACC | CCACAGTTGGTGGCATCTCG |
| *CG10725* | GATCGCTGTGACTATCCGCA | GCAGTTGACTTTCGAGGGGA |
| *CG4734* | TCACCATTGAGAGTCGTGCC | TAGTATCGGTTTGGCGACGG |
| *Sgs7* | GCAGTCACCATCATCGCTTG | ATCGCTAATGAGCTGCTGACA |
| *CG9682* | ATCAAGTGTCCCAGGAAGCG | TGCAAGCACAGGTAACCGAA |
| *CG10154* | TATTACGGCAAACCCGTGCT | TGGCCACACAGTCCACATAC |
| *Cda9* | TCCAGATGGTTGTGGAGCCG | TCTCGCTCGATGGTCGGAAC |
| *Muc55B* | AACCGCATCAGTGCCAAGTA | GAGTAGTGACCGTGACTGGC |
| *Sgs8* | ATTGCGTGCATCATGCTCATC | CCGCTCAAGACCCTCCATAA |
| *CG6283* | GAGTCCATCCAGACCAACGG | GCCTCGGCGTAGTAGAGAAC |
| *LysC* | CGACCAGCTGAACAAGTGAG | GTAGTCGTTGGAGCCGTTGTA |
| *thetaTry* | AGATACTGTTGTCACGGCGG | AGCTTCAGTATGCCCACGTC |
| *epsilonTry* | CAGTCGATTGAGGCCAAGGA | AGGGTTGGAGTCGGCAATAC |
| *CG32302* | AGAGGTCCCTTCAGCTGTCA | AGAAGTACCGATTGTCGGGC |
| *Vha100-4* | AAGGTGTGCACTGGTTTCCA | GTCCTGTGGTCACTCGTCTG |
| *LysD* | ACAACGAATGCGGATTGAGC | CAGTCATCGATGGACGGCAA |
| *CG17191* | GGCAGTTTCGAGTGGATGGA | ATGCCATCAGTGTAGCGACC |
| *obst-F* | GCCGTCACTATGGAGCCTAC | CACTCAACTGGCACTCGGAT |
| *CG12934* | CCACAGCACCACAACTCTCT | GCTTGACGCCCAAAGAAACC |
| *CG33128* | ATCGACGAGCCAGTCTTTGG | CACGGTTCCATTCCAGGTGA |
| *mt:srRNA* | GGCGGTATTTTAGTCTATCTAGAGG | ACAAATTTAAGTAAGGTCCATCGTG |
| *Tsp42Ep* | AGCACCGCGATAAGCTCTAC | CCGCTGTCGTAGTCGAAAGA |
| *CG6403* | GCAAGGTGATAGCCGACGAA | GTTCACACAGGAGGGCTTCA |
| *CG7715* | ATTGTGCTTCTTGAGCCTGC | ATATAGTGGGCTGGCGACAC |
| *obst-J* | GCCGAGAAACAAGACCTGGA | CGGACCACTTCCTTTTCGGA |
| *CG34279* | AAATTGTCTGCACGTTGGGC | CACACAACTGCTGGCAAAGG |
| *LysS* | CCTCTTGACCGACGACATCA | CTGCAGTAATGCCAGACGGC |
| *CG6839* | TCAACAATGAACTGGGCGGA | CGGGAGCGTAGTCGAAATCA |
| *CG15153* | CAGGCCACTCACTACTGTGG | CACCAGGGAATCCTGTGACC |
| *CG44956* | ACCGCGAAGACTTCACTTCA | GCCTCTTGGCAAATCCATCC |
| *CG12715* | TACGATGGCGATAACCAGCC | ATTCGCTTTGGACGTTGTGC |
| *CG17105* | GCTATCCTGCTCCTTGCCA | AACCAACGCGTCCTCCTAAG |
| *CG43896* | GCTCCGACCAGCGATACTAC | GTAGTGCTGCTTCGGACAGT |
| *CG4563* | TTCCACGCCGTCAACTACTC | GATCTCCTTGATGCGGTCGT |
| *alphaTry* | CGTCTGAGCTCTTCCCTGAG | CCGTATCCGTAGGTGGAGGA |
| *LysB* | GGTCCCGAGAACTACAACGG | AGCTCAACCCGCACTCATTG |
| *Obp99b* | ATTGGGTCTGGCCTTTGTCC | CAGATCCTCGTGCGTCTTCA |
| *Lcp65Ac* | GAAGTGCACAGTTGCCATCG | ACGTTCTTGAGCTGACCCTG |
| *CG6277* | GAACAGTGGATGGAAGCCCA | AGCTGGAGGCATCAATGGAC |
| *Mal-A6* | ATGCCACTACCGGGAAAAGG | CGTTCATCGTTCCACTCCCA |
| *CG3819* | ATGCTTCAACGAGGACGAGG | CGCCGGCTCCAAATGTAATG |
| *CG5767* | CAACGCCAAGCCCAATGTTC | GGTGGTCACATCAGCGGAAA |
| *Cpr64Ad* | GGGACCTGCTCCATACTTCG | AGGGGCGTTGTATCCCAAAG |

| | | |
|---|---|---|
| *Jon25Bi* | GTCCAGCAGGTTCATCCCAA | CGTGGGCGATGATAGAACCA |
| *CG34026* | CGAAGGCGTTTTTCGTTGGT | CCTTGATGCCGTAACCTCGT |
| *TwdlP* | AGGCGTCAGTTCACAAGGAG | AAACGGCATTCTCGTAGCCA |
| *Lsp2* | CCAGGACGATGTGTTCACCA | GCTGGAAGGTCTCCCAGTTC |
| *CG8629* | TCTCCAAGAAGCCCACCGAC | AGGCGTACTTGGGGGCATAC |
| *CG11672* | ATTGCCGTAAGTGTCAGCGA | CCCAAGTCTTCCGTACACCC |
| *CG43139* | GCAACTGGTTTTGCATCCTGT | GAACGTCACAAGACCATCGG |
| *CG12115* | ACTAGACTTTCTGGGCTGCG | CAGGATGCCCTTGGTGGTAA |
| *deltaTry* | ATCGGTCTGGCTAGCTCCA | CCGTATCCGTAGGTGGAGGA |
| *CG4830* | ACTACTGGAATGCCCAAGGC | CCCACAGCCCAGTAATCCAG |
| *Pebp1* | AAGGAGCTGACTCCCACTCA | CCTTGTTGCCGGGAATGTTG |
| *Lcp65Af* | GGCTCTCAACGTCAAGGGTA | CGGTGTAGGCGATGGAGTAAG |
| *CG34251* | CAGAGTGTGCCAGTGTACGA | ACTGATCCCGTAGCCATCCA |
| *Mal-A1* | GATCCGAATGCGTGCAACTC | AAGATCTGCAGGTGGGAACG |
| *Alp4* | GCCACTCATACGCACCCTAA | CCTTGTAGATGCGACCAGCA |
| *CG42235* | GAGGGCGGACGCCTAAATAC | GACGCATCCCATGTAGAGCA |
| ***Phae1*** | AGTGCTCCCTATGCGGTGTC | GTGCTTGCAGTTCCGTCCAC |
| *CG10513* | AACCTACTTGGAGCGACTGC | GCGGGCTTGATCACCAAATC |
| *gammaTry* | ATCGGTCTGGCTAGCTCCA | CCGTATCCGTAGGTGGAGGA |
| *CG10140* | AATGCCAGGATGGCCTTCAA | CAGGTTTGCTCCCTGGGAAT |
| *CG30016* | ATACTTCGGTGGGAAAGGCG | TAATAGGCGCCCACGTGAAA |
| *betaTry* | CATCGCTGTGCTCCATCTGA | GGAGCCAGAGGACTCGGTA |
| *CR11538* | CTGCAGCTAACGCAATCACC | ATCAAGGCTACGGATGCGAC |
| *CG7587* | AGATCTACTGGCGCGAACAC | CATCCAAGGAAGCACTTGCG |
| *CG7606* | GTCCAGAGCTGTTCCGTTGA | CCGGAAGAGGAAGGACATCG |
| *CG10405* | CCCTTCTACGAACAGCGGAC | CCTGCTTACGCTTCGGTGA |
| *CG4363* | GAGGATCCATTGTGCGTGGA | GCAGTTGGCATCGTTCAGAC |
| *Mal-A8* | ACAAACGAAGAGCAGCCAGT | CAACCCCACGGATCAACAGT |
| *CG7017* | TGCTAATCTCGGCTACTGCG | CTGTCCAGGCAGAAGGTGTT |
| *CG32557* | GCAAGCGGATCGTTAGCAAG | AGCGGAATTTGGACTGGAGG |
| ***z*** | TTACCTTCAGCGCCC | CCTCCCCCATTACCT |
| ***rp49*** | GATCGTGAAGCGCACCAAG | CCGGATTCAAGAAGTTCCTGGTG |

**Table S3.** Fly stocks

| No. | STRAIN | SOURCE (#STOCK No.) |
|---|---|---|
| TM001 | *$w^{1118}$* | BDSC (#BDSC_5905) |
| TM002 | *Phae1[SK1]* | Kondo S(#M2L-1221) |
| TM003 | *Phae1[Df]/CyO* | BDSC (#BDSC_3225) |
| TM004 | *Phae1-T2A-GAL4* | This study |
| TM005 | *Phae1-$HA^{wildtype}$* | This study |
| TM006 | *Phae1-$HA^{mutant}$* | This study |
| TM007 | *UAS-Phae1* | This study |
| TM008 | *UAS-Phae1 RNAi_1* | VDRC (#105269) |
| TM009 | *UAS-Phae1 RNAi_2* | NIG-FLY (#16996R-3) |
| TM010 | *$z^{v77h}$* | Kyoto DGGR (#101276) |
| TM011 | *$z^{Ty1}$* | This study |
| TM012 | *UAS-z-CC* | FlyORF (#F003293) |
| TM013 | *UAS-z RNAi_1* | VDRC (#109831) |
| TM014 | *UAS-z RNAi_2* | NIG-FLY (#HMJ21949) |
| TM015 | *UAS-mTor RNAi* | BDSC (#BDSC_33951) |
| TM016 | *UAS-GFP* | Tsuzuki S(12) |
| TM017 | *hs-gal4* | Tsuzuki S(12) |
| TM018 | *elav-Gal4* | Kyoto DGGR (#105-921) |
| TM019 | *nSyb-GAL4* | BDSC (#BDSC_51941) |
| TM020 | *repo-GAL4* | BDSC (#BDSC_7415) |
| TM021 | *c765-GAL4* | BDSC (#BDSC_602908) |
| TM022 | *phtm-GAL4* | BDSC (#BDSC_80577) |
| TM023 | *cg-GAL4* | BDSC(#BDSC_7011) |
| TM024 | *Tkg-GAL4* | Miura M (13) |
| TM025 | *Myo1A-GAL4* | Akagi K (14) |
| TM026 | *Eip71CD-GAL4* | BDSC (#BDSC_6871) |
| TM027 | *UAS-Phae2 RNAi* | VDRC (#108773) |
| TM028 | *UAS- CG9682 RNAi* | VDRC (#106391) |
| TM029 | *UAS- CG10154 RNAi* | VDRC (#101133) |
| TM030 | *UAS-Cda9 RNAi* | VDRC (#37981) |
| TM031 | *UAS- LysC RNAi* | VDRC (#39183) |
| TM032 | *UAS- thetaTry RNAi* | VDRC (#4593) |
| TM033 | *UAS- epsilonTry RNAi* | VDRC (#30807) |
| TM034 | *UAS- Vha100-4 RNAi* | VDRC (#330495) |
| TM035 | *UAS- LysD RNAi* | VDRC (#49814) |
| TM036 | *UAS- obst-F RNAi* | VDRC (#106346) |
| TM037 | *UAS- mt:srRNA RNAi* | VDRC (#103272) |
| TM038 | *UAS- alphaTry RNAi* | VDRC (#103292) |
| TM039 | *UAS- Cpr64Ad RNAi* | VDRC (#4488) |
| TM040 | *UAS- Jon25Bi RNAi* | VDRC (#105042) |
| TM041 | *UAS- TwdlP RNAi* | VDRC (#46487) |
| TM042 | *UAS- Lsp2 RNAi* | VDRC (#14069) |
| TM043 | *UAS- CG4830 RNAi* | VDRC (#21633) |
| TM044 | *UAS- Pebp1 RNAi* | VDRC (#101957) |
| TM045 | *UAS- Lcp65Af RNAi* | VDRC (#23530) |
| TM046 | *UAS- Mal-A1 RNAi* | VDRC (#15789) |
| TM047 | *UAS- gammaTry RNAi* | VDRC (#31016) |
| TM048 | *UAS-betaTry RNAi* | VDRC (#50091) |
| TM049 | *UAS-Mal-A8 RNAi* | VDRC (#15798) |

**Dataset S1.** Data from the RNAseq, qPCR, and bioassay screens for stress-induced death regulators.

**Dataset S2.** Data from the chemical screen for the signaling pathways upstream of *Phae1* gene expression.

**Dataset S3.** Raw data (qPCR, bio-assay, TUNEL, Dcp-1 activation).
